# Dicentric chromosome breakage in *Drosophila melanogaster* is influenced by pericentric heterochromatin and reveals a novel class of fragile site

**DOI:** 10.1101/2022.10.13.512123

**Authors:** Hunter Hill, Danielle Bonser, Kent G. Golic

## Abstract

Chromosome breakage plays an important role in the evolution of karyotypes, and can produce deleterious effects within a single individual, such as aneuploidy or cancer. Forces that influence how and where chromosomes break are not well understood. In humans, breakage tends to occur in conserved hotspots called common fragile sites (CFS), especially during replication stress. By following the fate of dicentric chromosomes in *Drosophila melanogaster* we find that breakage under tension also tends to occur in specific hotspots. Our experimental approach was to induce sister chromatid exchange in a ring chromosome to generate a dicentric chromosome with a double chromatid bridge. In the following cell division, the dicentric bridges may break. We analyzed the breakage patterns of three different ring-*X* chromosomes. These chromosomes differ by the amount and quality of heterochromatin they carry as well as their genealogical history. For all three chromosomes, breakage occurs preferentially in several hotspots. Surprisingly, we found that the hotspot locations are not conserved between the three chromosomes: each displays a unique array of breakage hotspots. The lack of hotspot conservation, along with a lack of response to aphidicolin, suggests that these breakage sites are not entirely analogous to CFS and may reveal new mechanisms of chromosome fragility.. Additionally, the frequency of dicentric breakage and the durability of their spindle attachment varies significantly between the three chromosomes and is correlated with the origin of the centromere and the amount of pericentric heterochromatin they carry. We suggest that different centromere strengths could account for this.

## INTRODUCTION

Chromosomes of higher eukaryotes typically have a single localized centromere. Chromosome rearrangements that conjoin two centromeres present a serious problem for the normal segregation of chromosomes during meiotic or mitotic divisions (Sturtevant and Beadle 1936; McClintock 1939; Novitski 1952). When two centromeres on the same chromatid segregate to opposite poles the stretched chromatin may break, delivering chromosomes with broken ends to the daughter cells. As a result, these cells are likely to suffer segmental aneuploidy. However, if cells continue to divide they may experience further genome rearrangements from breakage-fusion-bridge cycles, recombination with other chromosomes, or chromothripsis (McClintock 1939; Ceccaldi *et al*. 2016). In *Drosophila*, cells with broken dicentric chromosomes most often die, either through the DNA damage response or as a consequence of aneuploidy (Titen and Golic 2008). Some cells with a broken chromosome may survive if they repair the broken end by telomere addition (healing), or by using the homologous chromosome as a template for repair (McClintock 1939; Haber & Thorburn 1984; Levis 1989; Biessmann *et al*. 1990; Malkova *et al*. 1996; Morrow *et al*. 1997; Ahmad and Golic 1998; Zhou *et al*. 2000; Gao *et al*. 2008; Malkova and Ira 2013; Bhandari *et al*. 2019). When dicentric chromosomes break in the germline, and are healed, they may be recovered in progeny. Broken and healed chromosomes *Y, 2, 3* and *4* have been recovered in this way (Ahmad and Golic 1998; Titen and Golic 2008; Titen and Golic 2010; Titen *et al*. 2014). However, for *X,* 2 and 3, healed chromosomes can typically be recovered only if the breaks are very near the telomeres, owing to lethal aneuploidy of chromosomes that lack more than approximately 1 Mb of euchromatin (Lindsley *et al*. 1972; Mason *et al*. 1984; Levis 1989; Biessmann *et al*. 1990; Ashburner *et al*. 2005; Ryder *et al*. 2007; Cook *et al*. 2010; Titen and Golic 2010). Thus, recovering the products of dicentric breakage within a long stretch of euchromatin has remained a challenge until recently.

As previously reported, we devised a scheme to recover broken-and-healed *X* chromosomes by inducing sister chromatid exchange (SCE) within a ring chromosome in mitotically dividing cells of the male germline (Hill and Golic 2015). This generates a dicentric chromosome with two chromatin bridges. If each bridge breaks, and the ends are healed, a linear chromosome is generated. If such a chromosome is near-euploid, it can be transmitted and recovered in progeny. The ring chromosome we used contains the entire euchromatic portion of the *X* chromosome and an approximately equal amount of heterochromatin, consisting largely of *Y-*chromosome derived sequence. If the two dicentric bridges were to break in random locations, then most viable linear chromosomes would be expected to have breaks in heterochromatin since there are no required genes in heterochromatin of this chromosome, and large duplications or deficiencies would be tolerated (**Figure 1**). Surprisingly, approximately half of the recovered linear chromosomes had breakpoints in euchromatin, and those breakpoints clustered predominantly into a limited number of breakage hotspots.

**Figure 1:**
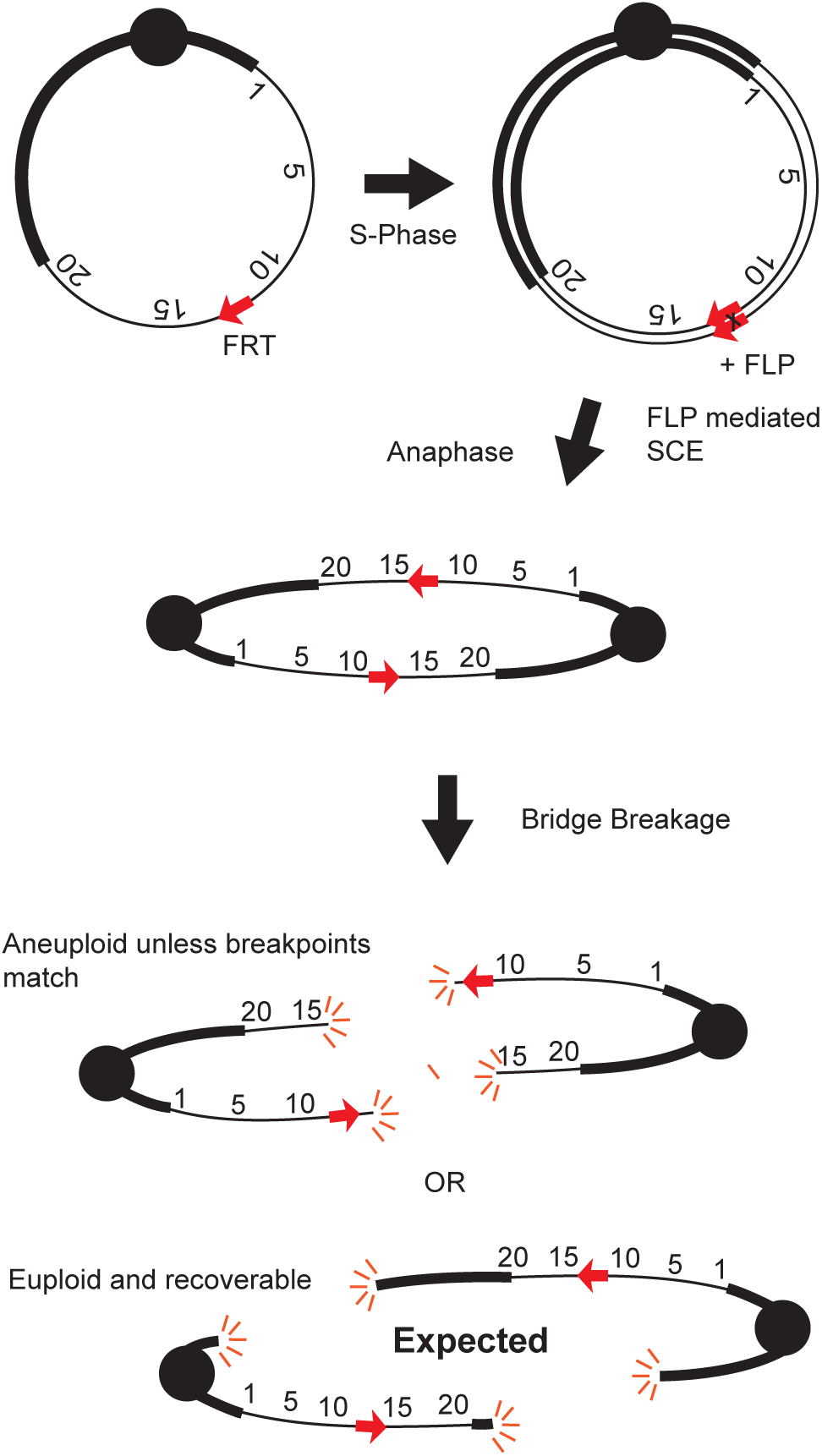
Dicentric bridges formed by sister chromatid exchange. FLP-mediated sister chromatid exchange within a ring generates double dicentric bridges, which can break in anaphase to yield linear chromosomes. Red arrows indicate *FRT*s; thick black lines indicate heterochromatin; thin black lines are euchromatin labeled with polytene map coordinates; black circles are centromeres. Top: breakage in euchromatin; bottom: breakage in heterochromatin.

Some of those breakage hotspots (cytological locations 11A and 13F) coincided with regions of the *Drosophila* genome that replicate late in cultured cells and are underreplicated in polytene chromosomes (Laird *et al*. 1987; Zhimulev and Belyaeva 2003; Belyaeva *et al*. 2008; Schwaiger *et al*. 2009; Yarosh and Spradling 2014). Late replication is also characteristic of common fragile sites (CFS) in the human genome (Le Beau *et al*. 1998; Hellman *et al*. 2000; Achkar *et al*. 2005; reviewed in Le Tallec *et al*. 2012). CFS appear as constrictions or breaks in metaphase chromosomes of cells experiencing replication stress, commonly induced by low concentrations of the DNA polymerase inhibitor aphidicolin (Glover *et al*. 1984). More recent work has shown that late-replicating CFS are often found far from active replication origins, confirming a connection to delayed replication (Palumbo *et al*. 2010; Brison *et al*. 2011; Le Tallec *et al*. 2011). Another interesting characteristic of CFS is that they reside in highly conserved chromosome locations throughout the entire human population, and in syntenic regions of mouse chromosomes (Helmrich *et al*. 2006). Chromosome breakage at CFS leads to carcinogenic genome rearrangements, including deletions of tumor suppressor genes (Hellman *et al*. 2002). In spite of this interest, human CFS have not been examined in the context of dicentric chromosomes, nor have the consequences of CFS breakage been examined in the germline.

In our previous work, not all dicentric breakage hotspots were correlated with sites of late replication. The present work was undertaken to more fully identify breakage hotspots in the ring chromosome used previously, and to similarly query other ring chromosomes to see if breakage hotspots were present, whether they were conserved, and to assess the effect of replication stress on breakage hotspots.

## MATERIALS AND METHODS

### Cytology

To observe metaphase chromosomes, the brains of third instar larvae were dissected in 0.7% NaCl, transferred to a hypotonic solution of 0.5% sodium citrate for 10’ at room temperature, and then fixed in a 11:11:1 solution of HOAc: MeOH: H2O. After fixation, the brain was squashed in 45% HOAc under a siliconized coverslip with bibulous paper and frozen on dry ice. The coverslip was flipped off with a razor blade and the slide air-dried. These squashes were stained with 4′,6-diamidino-2-phenylindole (DAPI) and viewed under UV fluorescence with a Zeiss Axioplan.

To examine anaphase figures, brains were fixed immediately after dissection, skipping the hypotonic incubation.

For polytene analysis of euchromatic breaks, third instar larvae were dissected in 45% acetic acid. Salivary glands were moved onto a coverslip with Lacto Aceto Orcein and squashed lightly. These spreads were analyzed on a Zeiss phase-contrast microscope.

### Recombination assay

When dicentric bridges were broken within heterochromatin, a recombination assay was performed to determine the location of the centromere relative to the orientation of euchromatin: *i.e.,* whether the chromosome was inverted or not. *R(1;Y)6AX2* and *R(1;Y)11AX2* are marked with *y*, so first we picked males with the opened ring and crossed them to *y^+^ w^bf^ f^5^* females. Next, we picked the heterozygous female offspring, crossed them to their brothers (5 vials of 2x2) and scored recombinant and non-recombinant offspring. A recombination frequency near zero indicates that the ring was broken to yield an inverted chromosome; normal levels of recombination indicate that the ring was broken to yield a linear-*X* in the normal orientation (*y* and *f* are separated by 56.7 map units)

*R(1)2* is marked by *f* as well as *y*, so we crossed males with *R(1)2* opened in heterochromatin to *y^+^ w^1118^*females, then crossed the heterozygous female offspring to their brothers, and scored their progeny as above. Recombination frequencies are found in Supplemental Material Files S1, S2, and S3.

### Somatic ring opening

To visualize somatic ring breakage, ring-*X* males were crossed with females homozygous for a heat shock inducible *FLP* transgene, (*70FLP*10, or *70FLP3F*) and allowed to mate at 25° until larvae were crawling on vial walls. To induce *70FLP*, vials were heat shocked for 1 hour in a water bath at 38° then removed from the bath and left at 25° for 6-8 hours or 16-18 hours. For anaphase preparations, vials were removed from the bath and left at 25° for 2-3 hours. Female third instar larval brains were dissected, and mitotic slides were prepared with DAPI as above.

### Feeding aphidicolin

Dry aphidicolin (338.49 g/mol) was ordered from MP Biomedicals [catalog # 159883] and 1 mg was diluted in 95% EtOH to 980uL to make a 3mM solution. Aliquots were further diluted with dH2O and manually stirred into vials containing warm liquid fly food to a final volume of 2 ml. Liquid fly food was then allowed to solidify at room temperature.

### Aphidicolin treatment of larval brains

Third instar larval brains were dissected and incubated in 1x PBS with 2.5 µM aphidicolin for 2-3 hours, and mitotic chromosomes were then examined via brain squash.

### Aphidicolin effect on fertility

We tested several different concentrations mixed in fly food to determine a dosage of aphidicolin that would have an affect without causing complete sterility. For this experiment, parents were allowed to lay eggs on fly food without aphidicolin for 24 hours. Embryos were counted and transferred to fly food with varied concentrations of aphidicolin. These flies were exposed to a consistent dose of aphidicolin for their entire development (embryo-adult). Finally, we mated each treated adult male with two untreated females and scored their fertility. Lower concentrations of aphidicolin (<50 μM) were shown to have no significant effect on the fertility of *Drosophila*. However, food that contains 100μM of aphidicolin has a significantly detrimental effect on fertility (P<0.01). These results prompted the use of 100 μM aphidicolin in subsequent experiments. This concentration has a clear detrimental effect while giving enough viability and fertility for successful experiments and substantial sample sizes. Additionally, wildtype larvae that are fed 100 µM aphidicolin in this fashion have chromosome constrictions and breaks similar to brains soaked in 2.5 µM aphidicolin.

### Acridine Orange

To visualize cell death after somatic dicentric breakage, ring-*X* males were crossed with *w^1118^; 70FLP10* females and allowed to mate at 25° until larvae were crawling on vial walls. To induce *70FLP*, vials were heat shocked for 1 hour in a water bath at 38° then removed from the bath and left at 25° for 24 hours.

For detecting apoptosis, wing disks were dissected from female third instar larvae in PBS. Both wing disks were immediately transferred to 5ug/ml acridine orange in PBS and allowed to soak for 2 minutes, they were then washed in PBS for 30 seconds before moving them to separate wells on a slide for visualization. A slide is prepared by adding a layer of electrical tape to the top, then cutting a small rectangle for each wing disk to sit in PBS under a coverslip. This allows for visualization without putting pressure on the tissue. Staining was observed by epifluorescence with UV excitation on a Zeiss Axioplan microscope using a 10X objective and intermediate 1.25X lens. All photos were taken with 480 ms exposure using a Zeiss Axiocam 702. Puncta within the large lobe of the wing disk were counted using Image-J using the ‘find maxima’ tool (example photo of area counted provided in Supplemental Material File S4).

### Statistics

For the Poisson distribution comparisons, each Chi-squared analysis was calculated using classes with at least 2 expected. For example, in the *R(1;Y)6AX2* polytene calculation with 19 bins: hits 0,1,2,3 were summed together as were 8, 9, 10, 11+ so that no categories had less than 2 expected. Degrees of freedom (df) were total categories minus two, so for this example there were six categories (4,5,6,7 and the two sums), therefore four df. The Chi-squared value was calculated using minitab software, and P values were obtained with an online calculator. (Supplemental Material File S5 & https://www.socscistatistics.com/pvalues/chidistribution.aspx)

For contingency tests if total N<300 then Fisher’s exact probability was used. If N>300 then Chi-square was used. For the gaussian distribution breakage comparison, the Anderson-Darling Normality test was used within Minitab software.

## RESULTS

### Collecting linear chromosomes from ring chromosome opening

FLP recombinase was used to induce SCE within a ring-*X* chromosome that carries *FLP Recombination Targets* (*FRTs*) at cytological location 13EF (**Figure 1**). FLP was expressed in mitotically dividing cells of the male germline using *nosGal4* and *UASFLP* transgenes, and *UAShiphop* to increase the rate of healing (Kurzhals *et al*. 2017). These males were mated to attached-*X* females, so that opened ring chromosomes would be recovered in sons. This scheme requires that the linear chromosome carry all vital *X*-linked genes to be recoverable. The attached-*X* females used in this cross also carried *eyFLP,* which expresses FLP in the developing eye (**Figure 2**). If a son inherits *eyFLP* and a ring-*X* chromosome, he will have small rough eyes as a result of ongoing dicentric formation and consequent cell death in the eye. However, if a son inherits *eyFLP* and a linear-*X* chromosome, his eyes will be normal (**Figure 2B,C**). To recover linearized *X* chromosomes we screened for *eyFLP-*bearing sons with normal eyes.

**Figure 2:**
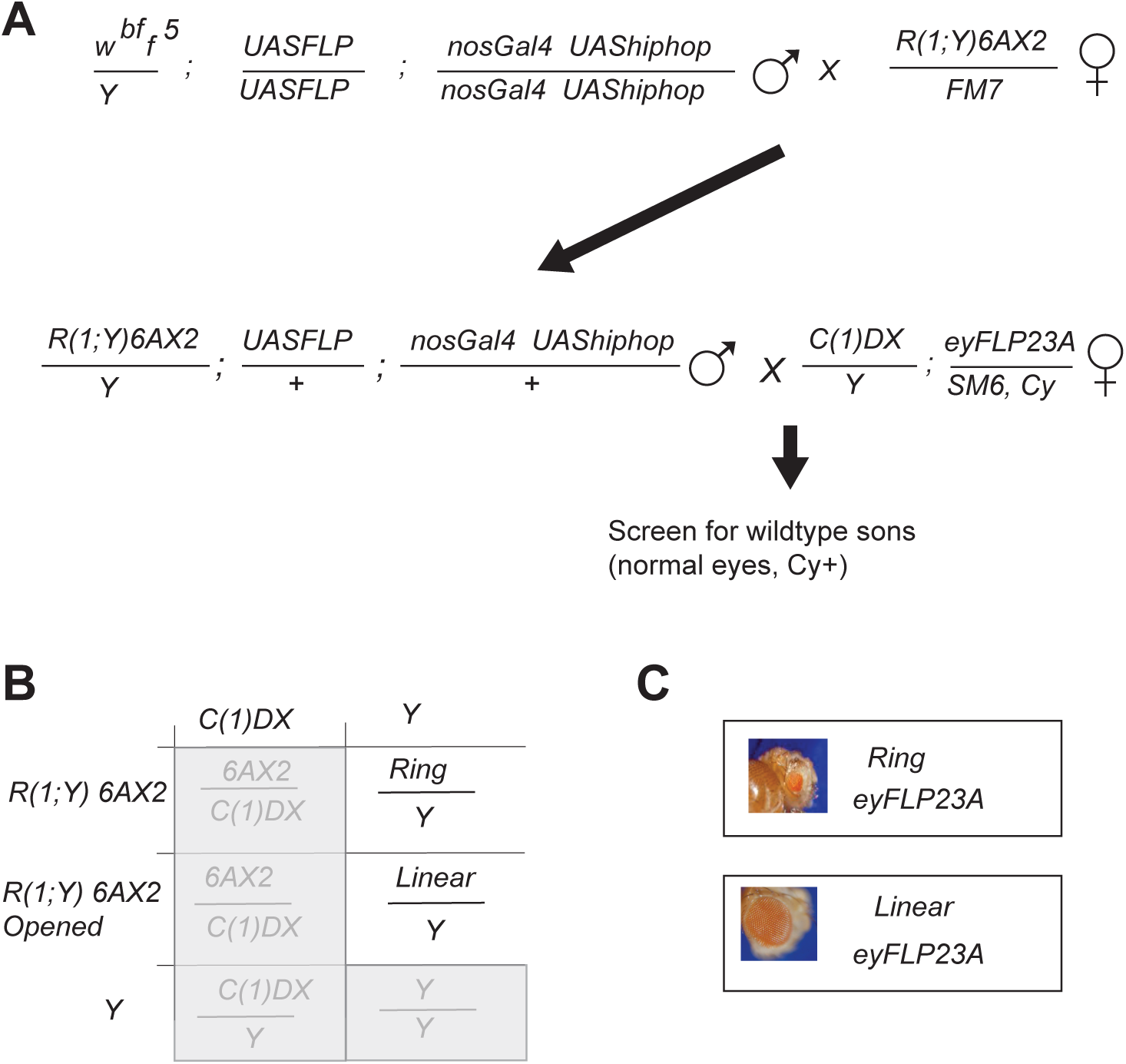
Genetic screen used to recover linearized chromosomes. A: Crossing scheme. B: Punnett square representing the final cross. Lethal and female genotypes are in grey. C: Eye phenotypes for viable males.

### Hotspots and coldspots of dicentric breakage

We previously isolated 25 openings of *R(1;Y)6AX2* and identified breakage hotspots in euchromatin in polytene divisions 11 and 13. To more completely identify hotspots (and coldspots) of dicentric breakage, we recovered 67 additional ring openings. The linearized chromosomes were characterized by mitotic cytology to confirm that they were linear and to make an initial determination of whether the chromosome termini were in euchromatin or heterochromatin (**Figure 3**; **Table 1**).

**Figure 3:**
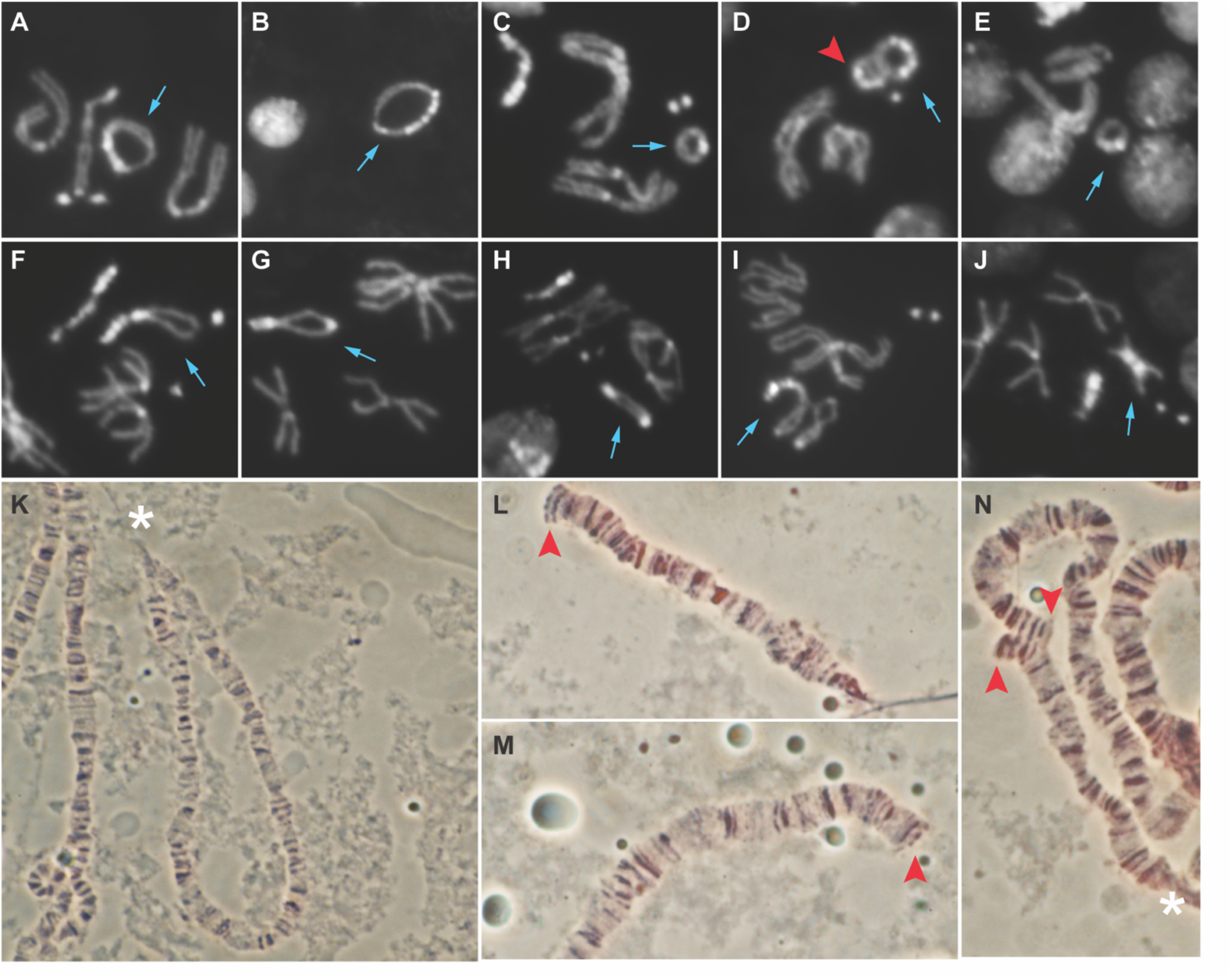
Cytology of ring chromosomes and opened rings. A-J: Mitotic Cytology A. *R(1;Y)6AX2/FM7*, with ring marked by blue arrow; B. *R(1;Y)11AX2*; C. *R(1)2*; D. *R(1;Y)6AX2/ R(1;Y)11AX2;* red arrowhead is *R(1;Y)6AX2,* blue arrow points to *R(1;Y)11AX2*. The rings appear to be interlocked in this nucleus; E. *R(1;Y)6AX2* opened in euchromatin with a large duplication that occasionally pairs to simulate a ring structure, as shown here; F. *R(1;Y)6AX2* opened in heterochromatin with very little heterochromatin at one end; G. *R(1;Y)6AX2* opened in heterochromatin with unequal, but obvious, heterochromatin at each end; H. *R(1;Y)6AX2* opened in heterochromatin with large blocks of heterochromatin at each end; I. *R(1;Y)11AX2* opened in heterochromatin; J. *R(1;Y)6AX2* opened in euchromatin; K-N. Polytene chromosomes from males with linearized rings. K. *R(1;Y)6AX2* broken in heterochromatin. Chromocenter marked by white asterisk; L,M. Linearized *R(1;Y)6AX2* (same as J) broken in euchromatin cytological locus 7B, with new telomeres marked by red arrowheads; N. *R(1;Y)6AX2* opened in euchromatin with a duplication spanning 12E-13D visible by pairing in polytenes (same as E) and new telomeres indicated by red arrows.

**Table 1:**
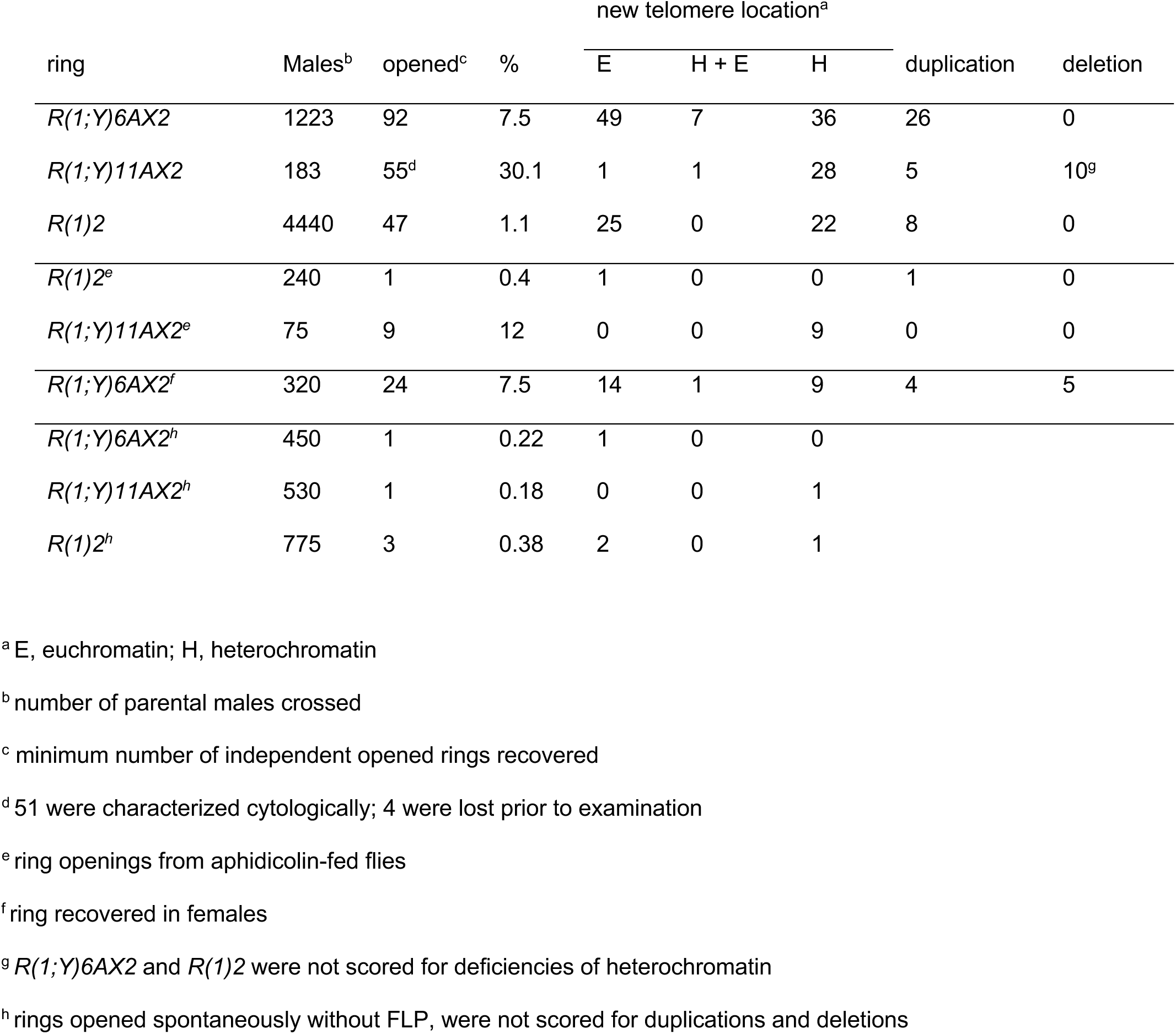
Germline ring openings

Chromosomes with breakpoints in heterochromatin could be identified either because they showed brightly staining blocks of heterochromatin on both ends of the chromosome, or because the termini of sister chromatids remain associated in metaphase spreads, a characteristic of heterochromatin (**Figure 3F-H**). In cases where there was any doubt, polytene chromosome analysis of males was also carried out to ascertain whether the chromosome formed a circular structure in salivary gland polytene chromosomes by virtue of both ends associating with the chromocenter (**Figure 3K**). Of the 67 new linear chromosomes, 25 were broken in heterochromatin, 38 were broken in euchromatin, and four were broken with one end in heterochromatin and one in euchromatin. With these 67 new linear chromosomes and 25 reported previously, a total of 92 linear chromosomes have been analyzed. The results described below refer to the aggregate results of these and the prior experiments.

To more precisely determine the locations of breakpoints in heterochromatin we first mapped the breakpoints relative to the centromere. A recombination assay was used to determine if the linearized chromosomes were normally oriented or inverted with respect to the centromere. Of the 43 chromosomes with at least one breakpoint in heterochromatin, 15 were oriented normally, with at least one break between the centromere and euchromatic division 1, while the other 28 were inverted, with at least one break between the centromere and division 20. Breaks in this latter, and larger, segment of heterochromatin could be further resolved: 11 of these 28 chromosomes were broken near the boundary of heterochromatin and division 20 of euchromatin, with DAPI-bright bands at only one end of the chromosome (**Figure 3F**); 13 had a large amount of DAPI-bright heterochromatin at one end but a only small amount at the opposite end (**Figure 3G**); while four had approximately equal amounts of DAPI-bright heterochromatin at each end (**Figure 3H**). These results show that chromosomes can break at several different locations in heterochromatin, and this distribution appears to be non-random. However, the limited resolution provided by mitotic chromosome cytology precludes drawing detailed conclusions about hotspots of breakage within heterochromatin.

For chromosomes opened in euchromatin, polytene chromosome analysis could be used to define the endpoints of the newly linear chromosomes. The distribution of breakpoints for the complete data set (25 + 67) shows openings occurring more often by breakage in euchromatin (105 chromosome termini) than in heterochromatin (79 chromosome termini), even though the heterochromatin and euchromatin regions are approximately the same size in metaphase (P *<* 0.005, assuming equal probability of breaks in heterochromatin and euchromatin). Only 103 breakpoints are mapped in **Figure 4A** because one stock with breaks in euchromatin was lost prior to polytene mapping. The breakpoints in euchromatin show clear hotspots and coldspots. When examined at higher resolution, hotspots are present even within a single numbered division (**Figure 5**).

**Figure 4:**
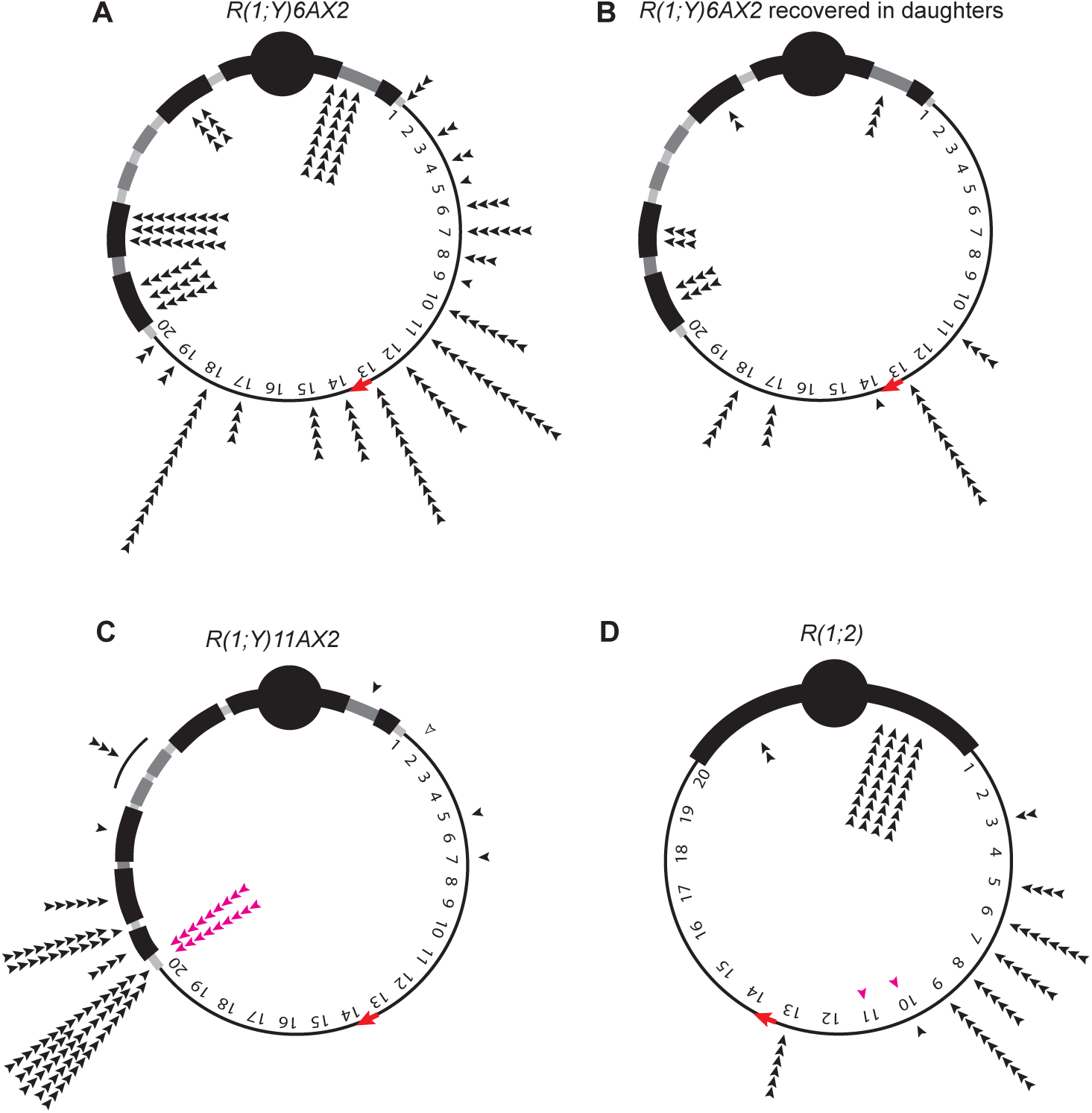
Breakpoint distributions for three different ring chromosomes opened in the male germline. Openings in euchromatin are indicated by polytene chromosome division; heterochromatin is divided according to DAPI staining blocks (Gatti and Pimpinelli 1983). Centromeres are represented by solid circles and their locations are estimated based on available data (Golic and Golic 2011; Ferree *et al*. 2014). Each arrowhead represents the location of a new telomere (two for each linear chromosome recovered; exact locations in supplemental material Files S1, S2, and S3; heterochromatin locations are approximate; open arrowhead indicates that location is unknown). A. *R(1;Y)6AX2* openings recovered in sons; B. *R(1;Y)6AX2* openings recovered in daughters; C. *R(1;Y)11AX2* openings recovered in sons; pink arrows mark chromosomes opened in the presence of aphidicolin; D; *R(1)2* openings recovered in sons.

**Figure 5:**
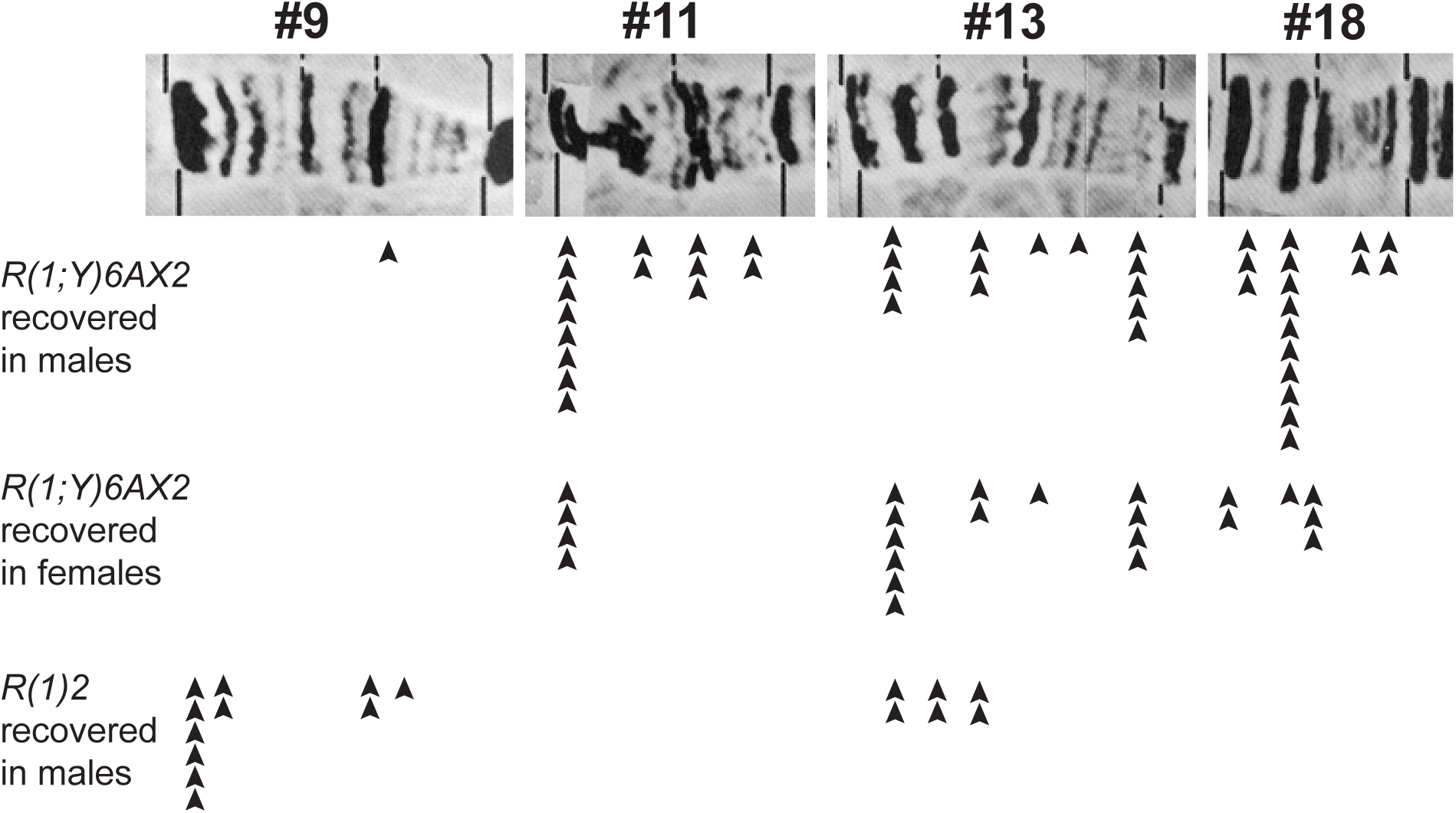
Close-up view of breakpoint distribution for four breakage hotspots of *R(1;Y)6AX2* openings recovered in sons, *R(1;Y)6AX2* openings recovered in daughters, and *R(1)2* openings recovered in sons. Segments of the Lefevre polytene maps (Lefevre 1976) are shown with breakpoints in regions 9, 11, 13, and 18. Each arrow indicates the location of a new telomere.

As demonstrated previously, divisions 11 and 13 of the polytene chromosome map are hotspots of breakage, but with this larger data set, division 18 has more breaks than any other polytene region. These three divisions (11, 13, 18) contain almost half (46/105) of all the breakpoints found in euchromatin. Two divisions, 2 and 16, were completely devoid of breakpoints, and several others (3, 4, 5, 9, 19, 20) had only one or two breakpoints.

Duplications are common among chromosomes opened in euchromatin: 46% (26/56) of chromosomes with euchromatic breaks have a duplication visible in polytene chromosomes. Some of these duplications are large enough to promote pairing of the chromosome ends in mitotic cells, where they sometimes appear as an intact ring (**Figure 3E,N**). However, such pairing only occurs in a fraction of nuclei, so linear chromosomes are still readily identified. The largest duplications contain more than a megabase of duplicated material (**Figure 6**).

**Figure 6:**
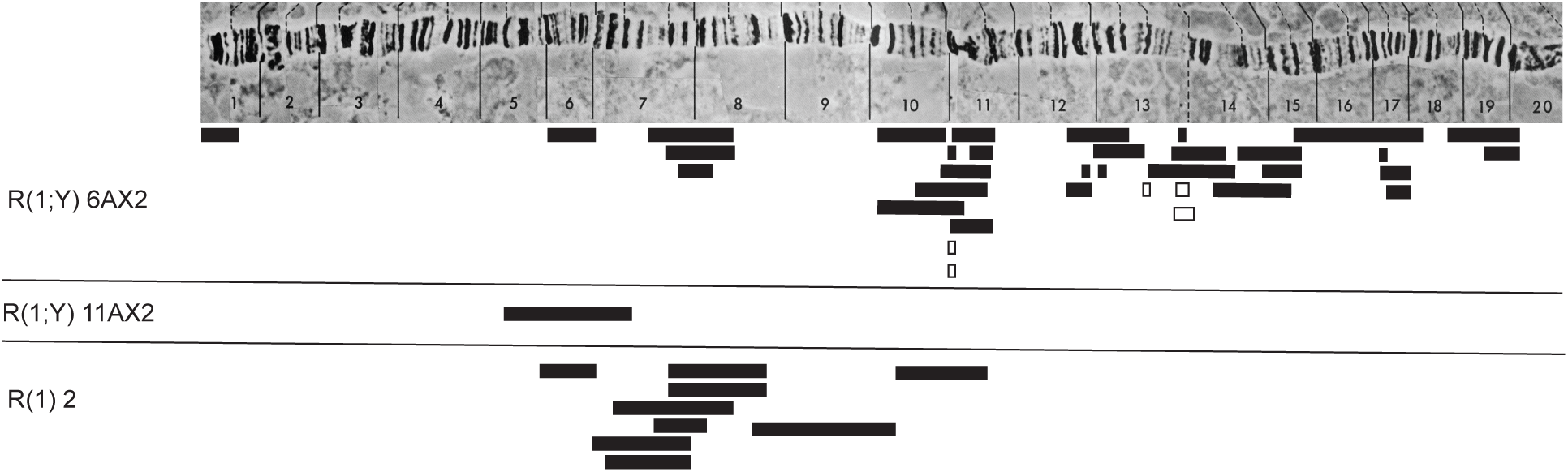
Duplications and deficiencies in the recovered linear chromosomes. The Lefevre photo map of the *X* chromosome is shown at the top. The extents of recovered duplications are indicated by solid bars; deficiencies (recovered in daughters) are shown as open bars.

We asked whether the distribution of breakpoints within euchromatin approximated a random distribution of breakpoints across all 20 numbered divisions by comparing our results to those predicted by a Poisson distribution. A Chi-squared goodness of fit test gives P < 0.05, indicating that they are not randomly distributed in polytene numbered divisions (**Table 2** left). However, not all polytene divisions carry equal portions of the genome — they vary up to ∼2X in their content. For instance, polytene division 11 accounts for ∼1.5 Mb of the *X* chromosome, while division 17 has only ∼0.8 Mb. This correlates with a large number of breakpoints in division 11 and a much smaller number in division 17. On the other hand, division 2 and division 18 both carry ∼0.96 Mb of *X* chromosome DNA, yet there were no breakpoints in division 2, and 17 breakpoints in division 18. To further test whether breakpoint hotspots are merely an artifact caused by the unequal distribution of DNA within polytene divisions, we re-plotted breakpoints into 1 Mb windows of the *X* chromosome using information from CytoSearch on flybase (http://flybase.org/cytosearch; File S6, **Table 1**). First, we searched the database for each whole number cytological location. Then, using gene sequence coordinates, we were able to identify the polytene subdivisions that correspond to whole number changes in megabases. For instance, ∼1 Mb of DNA exists between cytological regions 1A-1E and three breakpoints fall in that region. Within the 22 divisions of 1 Mb each in *X* euchromatin, hot and coldspots are still present, and the results do not fit the Poisson expectation (P < 0.01, **Table 2** middle). Therefore, even when breakpoints are allocated to equal size segments, they still show non-random clustering.

**Table 2:** Poisson prediction and goodness of fit to determine if break distribution is random

| | $R(1;Y)6AX2^a$ | | | | $R(1;Y)6AX2^b$ | | | | X-Ray | | | |
| --- | --- | --- | --- | --- | --- | --- | --- | --- | --- | --- | --- | --- |
|  | 101 breaks |  |  |  | 101 breaks |  |  |  | 133 breaks |  |  |  |
|  | 19 bins |  |  |  | 22 bins |  |  |  | 19 bins |  |  |  |
| | $\chi^2$ sum = 10.8512 | | | | $\chi^2$ sum = 15.789 | | | | $\chi^2$ sum = 1.97005 | | | |
|  | DF = 4 |  |  |  | DF = 5 |  |  |  | DF = 4 |  |  |  |
|  | P = 0.0283 |  |  |  | P = 0.0075 |  |  |  | P=0.7413 |  |  |  |
| Hits | p | Exp | Obs | $\chi^2$ | p | Exp | Obs | $\chi^2$ | p | Exp | Obs | $\chi^2$ |
| 0 | 0.00491 | 0.09329 | 2 | 38.9703 | 0.00912 | 0.20064 | 4 | 71.9455 | 0.00091 | 0.01729 | 0 | 0.01729 |
| 1 | 0.02612 | 0.49628 | 1 | 0.5113 | 0.04283 | 0.94226 | 1 | 0.0035 | 0.00638 | 0.12122 | 1 | 6.37068 |
| 2 | 0.06942 | 1.31898 | 4 | 5.4496 | 0.10059 | 2.21298 | 4 | 1.443 | 0.02234 | 0.42446 | 1 | 0.78039 |
| 3 | 0.12301 | 2.33719 | 2 | 0.0486 | 0.15752 | 3.46544 | 3 | 0.0625 | 0.05212 | 0.99028 | 1 | 0.0001 |
| 4 | 0.16347 | 3.10593 | 2 | 0.3938 | 0.18499 | 4.06978 | 3 | 0.2812 | 0.09122 | 1.73318 | 1 | 0.31015 |
| 5 | 0.17380 | 3.3022 | 1 | 1.605 | 0.17380 | 3.8236 | 0 | 3.8236 | 0.12771 | 2.42649 | 3 | 0.13555 |
| 6 | 0.15398 | 2.92562 | 2 | 0.2929 | 0.13608 | 2.99376 | 2 | 0.3299 | 0.14900 | 2.83100 | 1 | 1.18423 |
| 7 | 0.11693 | 2.22167 | 0 | 2.2217 | 0.09132 | 2.00904 | 1 | 0.5068 | 0.14900 | 2.83100 | 2 | 0.24393 |
| 8 | 0.07770 | 1.4763 | 2 | 0.1858 | 0.05363 | 1.17986 | 1 | 0.0274 | 0.13037 | 2.47703 | 3 | 0.11041 |
| 9 | 0.04589 | 0.87191 | 0 | 0.8719 | 0.02799 | 0.61578 | 0 | 0.6158 | 0.10140 | 1.9266 | 2 | 0.00280 |
| 10 | 0.02439 | 0.46341 | 0 | 0.4634 | 0.01315 | 0.2893 | 0 | 0.2893 | 0.07098 | 1.34862 | 1 | 0.09012 |
| 11+ | 0.02038 | 0.38722 | 3 | 17.6298 | 0.00899 | 0.19756 | 3 | 39.7533 | 0.09857 | 1.87283 | 3 | 0.67839 |

p = Poisson predicted probability
Exp = expected number of bins to contain X hits
Obs = Observed number of bins to contain X hits
$\chi^2 = (Obs - Exp)^2 / Exp$
a= bins are polytene regions 1-19
b= bins are 1MB each

### The coincidence of similar breakpoints at each end of linearized chromosomes

When a dicentric bridge breaks, we see no obvious reason why it should break particularly in heterochromatin or euchromatin. From prior work we know that breaks can occur both in heterochromatin and euchromatin, and that breaks in either location are capable of being healed (Ahmad and Golic 1998; Titen and Golic 2010). However, our scheme for recovery of linearized ring chromosomes imposes the additional requirement that the chromosome be viable and fertile in males. Since vital genes are spread throughout *X* euchromatin, only very small deficiencies of euchromatin will be tolerated. Duplications of euchromatin may be substantially larger, up to ∼10% of *X* euchromatin, though they typically exhibit reduced viability (Cook *et al*. 2010). *A priori*, it is expected that randomly occurring breaks in each of the twin bridges formed by SCE in a ring would rarely match closely enough in euchromatin to produce viable combinations. In contrast, the heterochromatin of this ring-*X* constitutes approximately half the chromosome and carries no genes that are needed for viability. Breaks in heterochromatin need not be closely matched for the chromosome to be recoverable, therefore it is much more likely that random pairs of breaks compatible with viability will be found in heterochromatin than in euchromatin. However, if breaks are not random but tend to occur at a limited number of hotspots, then the probability of viable two-break combinations in euchromatin is increased. Our results, which identify several hotspots, suggest that this is true.

An alternative explanation is that the breaks are not, in fact, independent, but that breakage and healing occurred spontaneously prior to S phase in an unreplicated chromosome. Matching ends would then be expected. If the site of breakage did not disrupt a vital gene, such chromosomes should be recoverable. However, such an event cannot easily explain the fact that the majority of breaks in euchromatin were found in chromosomes with duplications. These chromosomes are much more easily explained by regional hotspots of breakage, where breaks in two bridges occur independently at sites that are close, but not identical.

If duplications are produced by two independent breaks, then reciprocal events with small deficiencies should also be produced. These would not be recovered in males where all vital genes must be present, but they might be recovered in females when heterozygous with a complete *X* chromosome. To test this we crossed the *R(1;Y); UASHiphop/+; nosGal4 UASFLP/+* males to *N/FM7, B; eyFLP/SM6,Cy* females. Linearized chromosomes were recovered in FM7 Cy^+^ daughters, and recognized as flies with eyes that showed the normal B/+ phenotype. If such daughters still carried a ring chromosome the eyes would be much smaller and rougher than typical B/+ eyes. We recovered 24 opened rings from this cross (along with 3 false positives, which were discarded). Five of 24 chromosomes were lethal in males. Polytene analysis suggests that all 5 of these chromosomes carried deficiencies (**Figure 6**, open boxes). These chromosomes also exhibited breakage hotspots, just as those recovered in males did (**Figure 4B**; **Table 1**).

### Hotspot conservation, fragile sites and replication stress

Dicentric breakage hotspots could be a manifestation in *Drosophila* of what are called Common Fragile Sites (CFS) in humans. CFS typically cover broad chromosomal regions of up to 100’s of kb, are responsive to replication stress, are highly conserved, and are frequently involved in chromosomal rearrangements (Glover *et al*. 1984; Hellman *et al*. 2002; Helmrich *et al*. 2006). Breakage hotspots in divisions 11 and 13 are correlated with late or under-replication of these regions in a variety of cells (Zhimulev and Belyaeva 2003; Nordman *et al*. 2011; Belyaeva *et al*. 2012; Yarosh and Spradling 2014) However, the division with the most breakpoints, division 18, is not a region showing late replication in these studies. This raises the question of whether dicentric breakage hotspots can be explained as analogs of CFS, or whether there are other causes for the breakage hotspots. To further examine the possibility that dicentric breakage hotspots are analogous to CFS, we experimentally examined two properties: conservation and response to replication stress.

### Breakage of ring chromosomes with different histories

We reasoned that if breakage hotspots were analogous to CFS in humans then they should be conserved between different ring chromosomes. *R(1)2*, was recovered decades ago as a detachment of *C(1)RM* (Morgan 1926; Morgan 1933; Schultz and Catcheside 1938). It carries a relatively small amount of heterochromatin, all seemingly derived from the *X*. The *FRT*-bearing *P{>w^hs^>}75B* transgene, which is on *R(1;Y)6AX2,* was crossed onto *R(1)2* to induce SCE at the identical site. *R(1;Y)11AX2* was generated in our laboratory by the previously published method, but differs in that it carries a large block of heterochromatin that is absent from *R(1;Y)6AX2*. This difference likely arose when the original ring chromosomes, which were highly filicidal, were irradiated to recover chromosomes with improved viability. It also carries *P{>w^hs^>}75B.* Both rings carry all vital euchromatic genes of the *X* chromosome and are viable in males as the only *X.* We then used the same screen as before to recover linearized chromosomes.

We recovered 47 openings of *R(1)2* and 55 openings of *R(1;Y)11AX2*. *R(1)2* opened very infrequently, while *R(1;Y)11AX2* opened at a relatively high rate (**Table 1**). In each case we identified breakage hotspots, but the distribution of breakage hotspots was very different for each chromosome. Like *R(1;Y)6AX2*, the majority of breaks in *R(1)2* were in euchromatin; but unlike *R(1;Y)6AX2*, where most breaks were found in what is normally the proximal portion of euchromatin (divisions 11-20), with *R(1)2* most breaks were found in the distal portion of euchromatin (divisions 1-10). There is some overlap between *R(1;Y)6AX2* and *R(1)2* hotspots, but *R(1)*2 had no breakpoints in divisions 11 or 18, which were the two most frequent regions of breakage with *R(1;Y)6AX2*, and *R(1;Y)6AX2* had only a single break in division 9, which was the most frequent site of breakage in *R(1)2* (**Figure 4**). *R(1;Y)11AX2* showed a quite different pattern of openings from the other rings with nearly all of its breaks concentrated around a DAPI-bright heterochromatic block near the junction of heterochromatin and euchromatin (**Figure 4C**). As was the case with *R(1;Y)6AX2,* several of the *R(1)2* and *R(1;Y)11AX2* openings carried duplications of material at their termini. Because there are no essential genes in heterochromatin, linearized *R(1;Y)11AX2* chromosomes that were deficient for portions of heterochromatin were also found.

The very different distributions of breakage hotspots with these three ring chromosomes argue strongly against the proposition that dicentric breakage hotspots are analogous to CFS in human chromosomes, at least with respect to conservation of their location.

### Replication stress and dicentric breakage

Fragile sites were originally identified in cells experiencing replication stress (Hecht *et al*. 1980; Glover 1981; Hecht *et al*. 1981; Glover *et al*. 1984). Replication stress can occur in any organism when complications arise during pre-replication, initiation, or elongation, resulting in a local delay in DNA synthesis (Reviewed in Gelot *et al*. 2015). To further explore whether there is any connection between CFS and dicentric breakage hotspots, we examined whether replication stress could increase the frequency of dicentric breakage or alter the array of breakage hotspots. First, we confirmed the effect of the DNA polymerase inhibitor aphidicolin on the production of fragile sites in somatic cells. Larval brains were dissected and incubated in 1X PBS with 2.5 µM aphidicolin for 2-3 hours, and mitotic chromosomes were then examined. We observed novel constrictions and chromosome breaks after exposure to aphidicolin (**Figure 7A-C**, **Table 3**).

**Figure 7:**
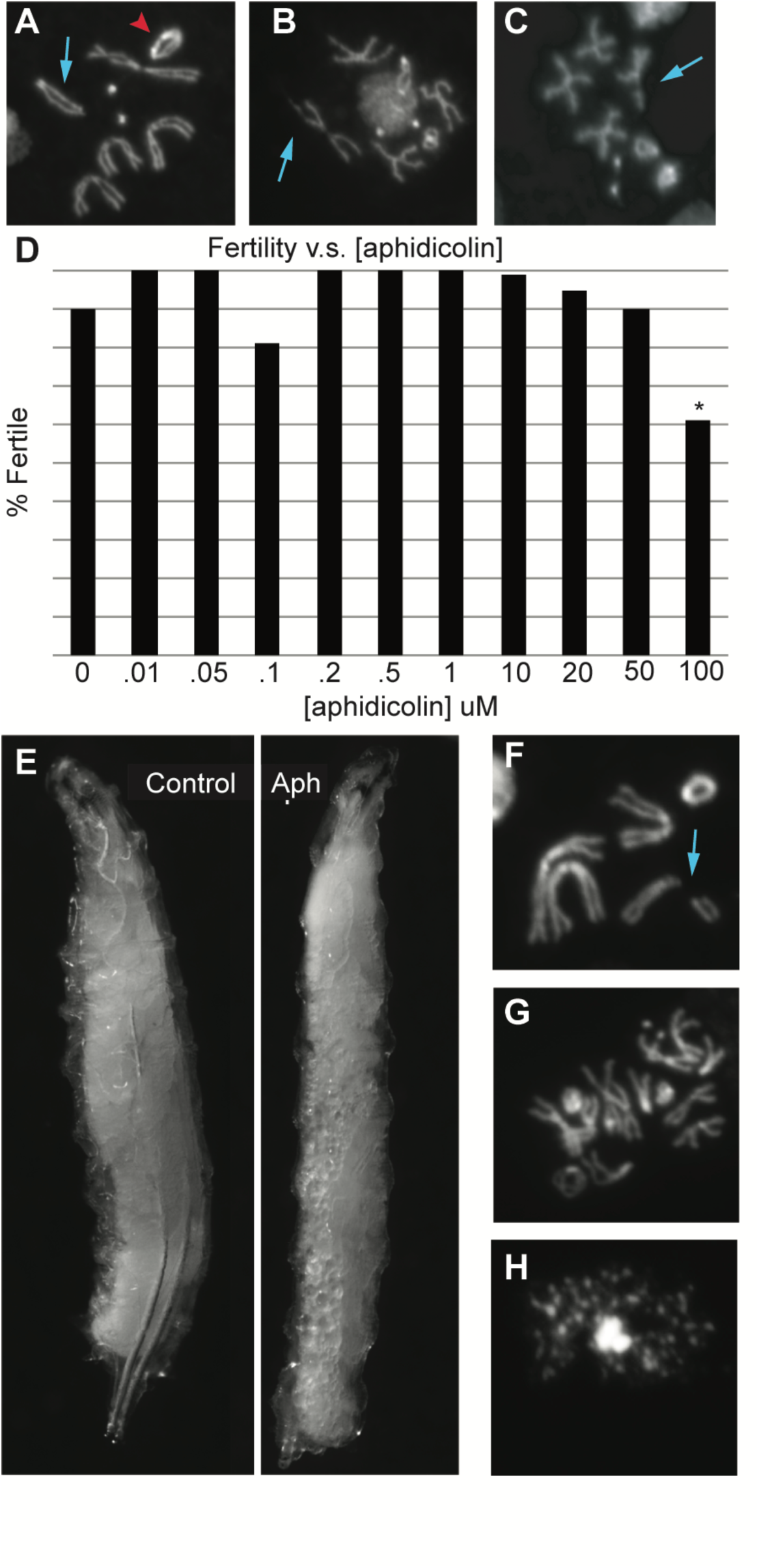
The effect of aphidicolin on chromosome breakage. A. Metaphase figure from female carrying *R(1;Y)6AX2/FM7*; red arrowhead indicates the ring; blue arrow points to the *FM7 X* chromosome balancer. B,C. Chromatid breaks appearing after soaking larval brains in aphidicolin; D. Dose/fertility results for flies reared on food containing aphidicolin (100 µM is only significant change in fertility, P=0.00047 2-tailed t test). E. Appearance of 3^rd^ instar larvae raised on normal food or food containing 100 µM aphidicolin; F. Chromosome break from larva raised on aphidicolin food; G. Polyploid cell from larva raised on aphidicolin food; H. Chromoanagenesis from larva raised on aphidicolin food.

**Table 3:** Somatic nuclei showing breakage after incubation in aphidicolin^a^

|  | number of breaks |  |  | % nuclei with breaks |
| --- | --- | --- | --- | --- |
|  | 0 | 1 | 2+ |  |
| - aphidicolin | 68 | 2 | 0 | 2.86 |
| + aphidicolin | 138 | 17 | 11 | 16.87<sup>b</sup> |

<sup>a</sup> *R(1;Y)6AX2/FM7* larval brains were dissected and incubated in 2.5 μM aphidicolin for 2-3 hours prior to fixation.
<sup>b</sup> P = 0.011, 3x2 contingency test

Next, we asked whether replication stress would affect the recovery of linearized ring chromosomes through the male germline. To administer aphidicolin to live animals we mixed aphidicolin into fly food to a final concentration of 100 µM, and raised flies from egg to adult on this medium. The decreased fertility of these flies proved the efficacy of aphidicolin in this system (**Figure 7D**). The larvae raised on aphidicolin were also delayed in development and had bodies that were filled with fat droplets (**Figure 7E**). Furthermore, when we examined neuroblast mitoses from these larvae we observed breakage at higher frequency than in the control, and polyploidy and chromoanagenesis were also observed (**Table 4**; **Figure 7F-H**) (Pellestor *et al*. 2022).

**Table 4:**
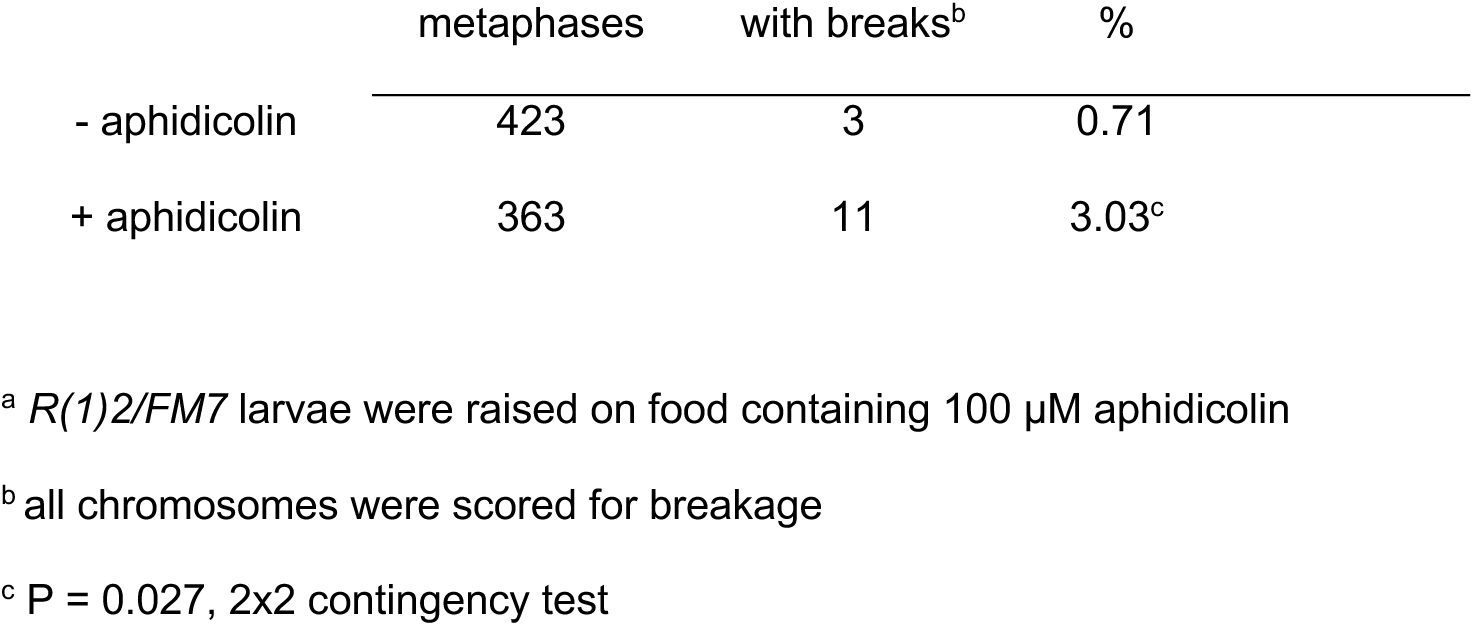
Somatic chromosome breakage from aphidicolin feeding of larvae^a^

|  | metaphases | with breaks<sup>b</sup> | % |
| --- | --- | --- | --- |
| - aphidicolin | 423 | 3 | 0.71 |
| + aphidicolin | 363 | 11 | 3.03<sup>c</sup> |

<sup>a</sup> *R(1)2/FM7* larvae were raised on food containing 100 μM aphidicolin
<sup>b</sup> all chromosomes were scored for breakage
<sup>c</sup> $P = 0.027$ , 2x2 contingency test

To analyze the effect of replication stress on dicentric breakage in the germline, we used *R(1;Y)11AX2* and *R(1)2*. *R(1;Y)11AX2* was chosen because there was a single hotspot of opening, and if new fragile sites were generated they would be quite obvious. *R(1)2* was chosen because it opened very infrequently, and if aphidicolin were to increase chromosome fragility, then the frequency of recovering linearized chromosomes might also be significantly increased. Males carrying one of these ring chromosomes, along with the same *nosGal4 UASFLP and UASHiphop* transgenes used previously, were fed aphidicolin and mated as before to recover linearized chromosomes. Aphidicolin did not change the distribution of recovered breaks with *R(1;Y)11AX2* (**Figure 4** pink arrows), nor did it significantly increase the rate of recovery of linearized chromosomes with either ring (**Table 1**). The linearized *R(1;Y)11AX2* chromosomes were all broken in the previously identified hotspot. *R(1)2* only yielded one opened ring when treated with aphidicolin; this chromosome had a duplication spanning the breakpoints in 10B and 11C. Duplications of this size were not uncommon when *R(1;Y)6AX2* was opened. Although *Drosophila* DNA replication is sensitive to aphidicolin, it does not have a noticeable effect on the recovery of linearized ring chromosomes through the male germline.

### Dicentric formation and chromosome breakage frequencies

In addition to striking differences in breakpoint distributions between the three ring chromosomes, the frequency of linear chromosome recovery varied greatly, from ∼1% of *R(1)2* males transmitting linear chromosomes to ∼30% with *R(1;Y)11AX2*. Since each of these ring chromosomes carries the same insertion of *FRT*s, and each experiment uses the same source of FLP, we expected the chromosomes to form dicentrics at very similar rates. Most males tested in our screens were sterile, indicating that all of their germline stem cells (GSCs) had experienced dicentric formation, but no viable healed linear chromosomes were produced. The high frequency of sterile males allows a determination of the rate of dicentric formation in the germline based on the number of false-positives, *i.e.,* fertile males that transmitted a chromosome which was still a ring. For *R(1;Y)11AX2* 183 males were tested and there were no false positives: every fertile male transmitted a linear *X* and the males which did not were sterile. Therefore the rate of dicentric formation for *R(1;Y)11AX2* was 100%. For *R(1;Y)6AX2,* 1223 males were tested and there were two false positives. Since each male carries ∼14 germline stem cells (Dansereau and Lasko 2008) ∼17,122 GSCs were screened giving a 99.99% cellular rate (17,120/17,122) of dicentric formation (making the assumption that, regardless of number of offspring produced by a particular male, each represents the clonal expansion of a single germline stem cell). With *R(1)2,* 4440 males (62,160 GSCs) were screened and there were 44 false positives, giving a rate of 99.93% dicentric formation. The rate of dicentric formation in the germline was at or very close to 100% in each case. Therefore, differential rates of dicentric formation cannot account for the differences in recovery of linear chromosomes (**Table 1**).

We next investigated whether dicentric chromosomes generated from different rings might break at different rates. For this, we used heat shock to induce the expression of a *70FLP* transgene in somatic cells. Ring-bearing males were crossed to females carrying *70FLP* and third instar larvae were then heat-shocked at 38° for one hour. Female larvae were dissected 1-2 hours after the end of heat shock and anaphase mitotic figures from brains were examined for bridges to determine the frequency of dicentric formation. Dicentric formation resulting from SCE was high in all cases, ranging from ∼91-100% with *70FLP3F* and ∼84-97% with *70FLP10* (**Table 5**).

**Table 5:** Dicentric anaphase bridges in larval brains after FLP induction

| FLP | Ring | Heatshock | Total | Normal | Bridges |
| --- | --- | --- | --- | --- | --- |
| 70FLP3F | *R(1;Y)6AX2* | - | 38 | 92.1% | 7.9% |
|  |  | + | 122 | 5.7% | 94.3% |
|  | *R(1;Y)11AX2* | - | 12 | 50% | 50% |
|  |  | + | 41 | 0% | 100% |
|  | *R(1)2* | - | 19 | 79% | 21% |
|  |  | + | 137 | 9.5% | 90.5% |
| 70FLP10 | *R(1;Y)6AX2* | - | 40 | 77.5% | 22.5% |
|  |  | + | 94 | 3.2% | 96.8% |
|  | *R(1;Y)11AX2* | - | 14 | 42.9% | 57.1% |
|  |  | + | 140 | 4.3% | 95.7% |
|  | *R(1)2* | - | 55 | 90.9% | 9.1% |
|  |  | + | 167 | 15.6% | 84.4% |

We then examined metaphase figures 6-8 hours after heat shock, giving cells time to divide with dicentric chromosomes and make it to the next metaphase. Cells were scored for whether they carried the ring chromosome or a linearized version. Both *R(1)2* and *R(1;Y)6AX2* showed approximately 5% of metaphases with broken *X* chromosomes, while *R(1;Y)11AX2* had ∼8% of cells with a linearized *X*. After normalizing to the rates of dicentric formation, *R(1;Y)11AX2* still produced linear chromosomes more frequently than the other chromosomes (8.6% for *R(1;Y)11AX2* vs. 4.9% for *R(1;Y)6AX2* and 5.9% for *R(1)2*). We examined metaphase figures again 16-18 hours after heat shock. At this time point there were pronounced differences, with *R(1)2* showing only 2.7% linearized rings, *R(1;Y)6AX2* showing 9.8% and *R(1;Y)11AX2* showing ∼20.5% (Figure 8).

**Figure 8:**
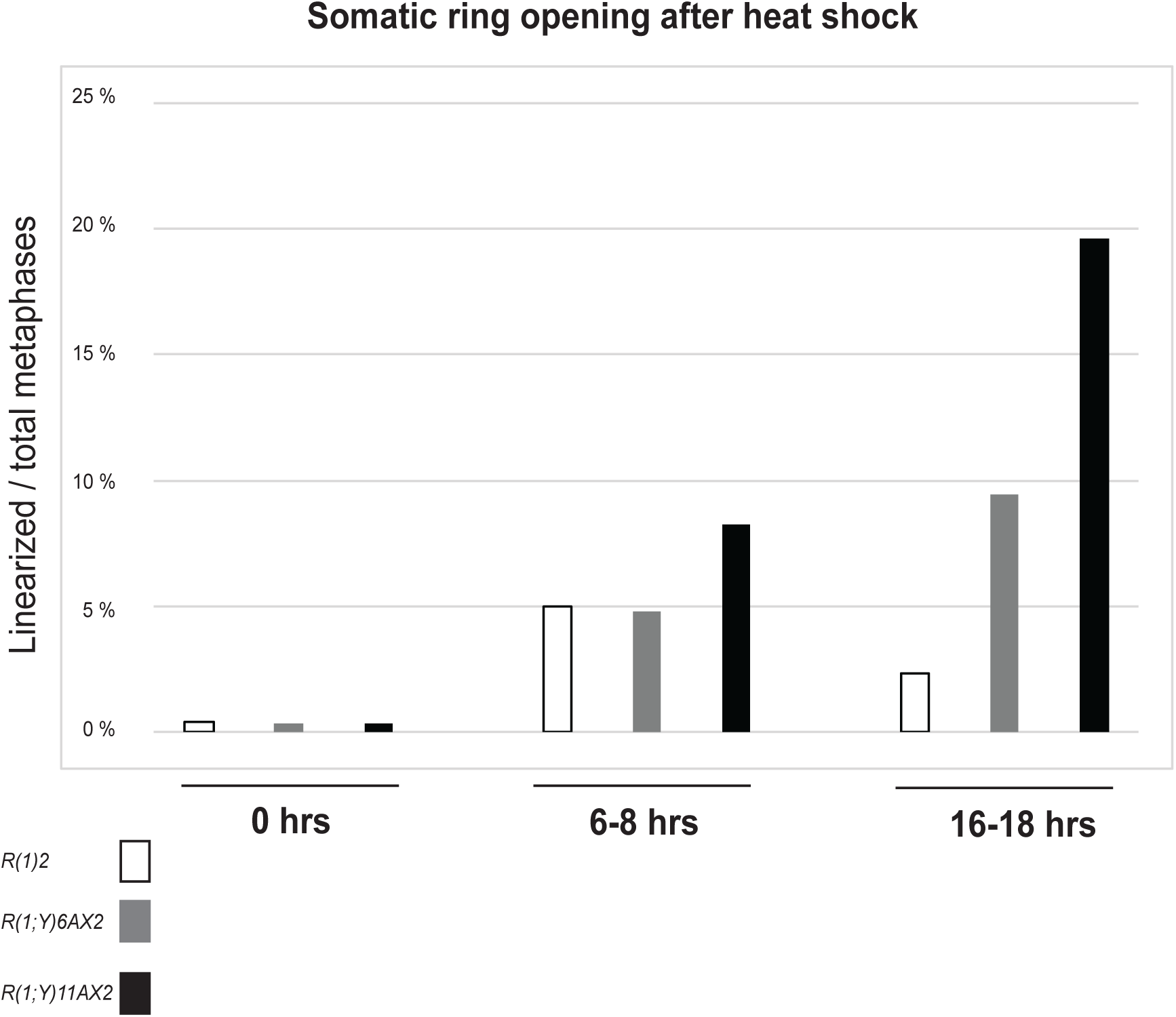
Histogram representing the fraction of somatic metaphases containing opened ring chromosomes after heatshock. A. *R(1;Y)6AX2,* N values represent total metaphases observed for each timepoint. N= 339, 567, and 201 for 0, 6-8, and 16-18 hours respectively; B. *R(1;Y)11AX2*, N= 640, 556, and 306; C. *R(1)2*, N= 267, 642, and 306.

At both time points, broken versions of *R(1;Y)11AX2* were clearly more prevalent than for *R(1;Y)6AX2* or *R(1)2.* The simplest interpretation is that *R(1;Y)11AX2* breaks most frequently and *R(1)2* breaks least frequently. This would readily account for the differences in their recovery rates through the germline.

Additional details of the cytological observations of anaphase bridges support the hypothesis that the rings break at different rates. While examining several hundred anaphases figures it became clear that chromosome bridges take several different forms. Many of the cells with dicentrics clearly showed the expected double bridge (**Figure 9B**). There were also anaphase/telophase figures giving the appearance of bridges that had broken (**Figure 9EF**). Surprisingly, we found that there were some anaphase figures with bridges, but instead of stretching and breaking, the dicentric lagged behind the rest of the chromosomes, seemingly released from the poleward spindle tension. (**Figure 9C&D**). We categorized these anaphases as “lagging”, and compared the frequency of lagging dicentrics observed with each of the three different ring chromosomes (**Table 6**). After heat shock induction of *70FLP3F*, *R(1)2* had the highest frequency of lagging dicentrics and *R(1;Y)11AX2* had the lowest. Conversely, *R(1)2* had the lowest frequency of broken bridges in anaphase, while *R(1;Y)11AX2* had the highest (3x3 contingency test comparing all three bridge categories, *P* = 0.0296). When the *70FLP10* transgene was used, *R(1)2* again had the highest frequency of lagging dicentrics and the lowest frequency of broken bridges (3x3 test, P = 0.0174). In this case *R(1;Y)11AX2* also had the lowest frequency of lagging dicentrics, but was very similar to *R(1;Y)6AX2* in the frequency of broken bridges. These results support the conclusion that differences in the frequencies of recovering linear chromosomes through the germline are at least partially dependent upon different breakage frequencies.

**Figure 9:**
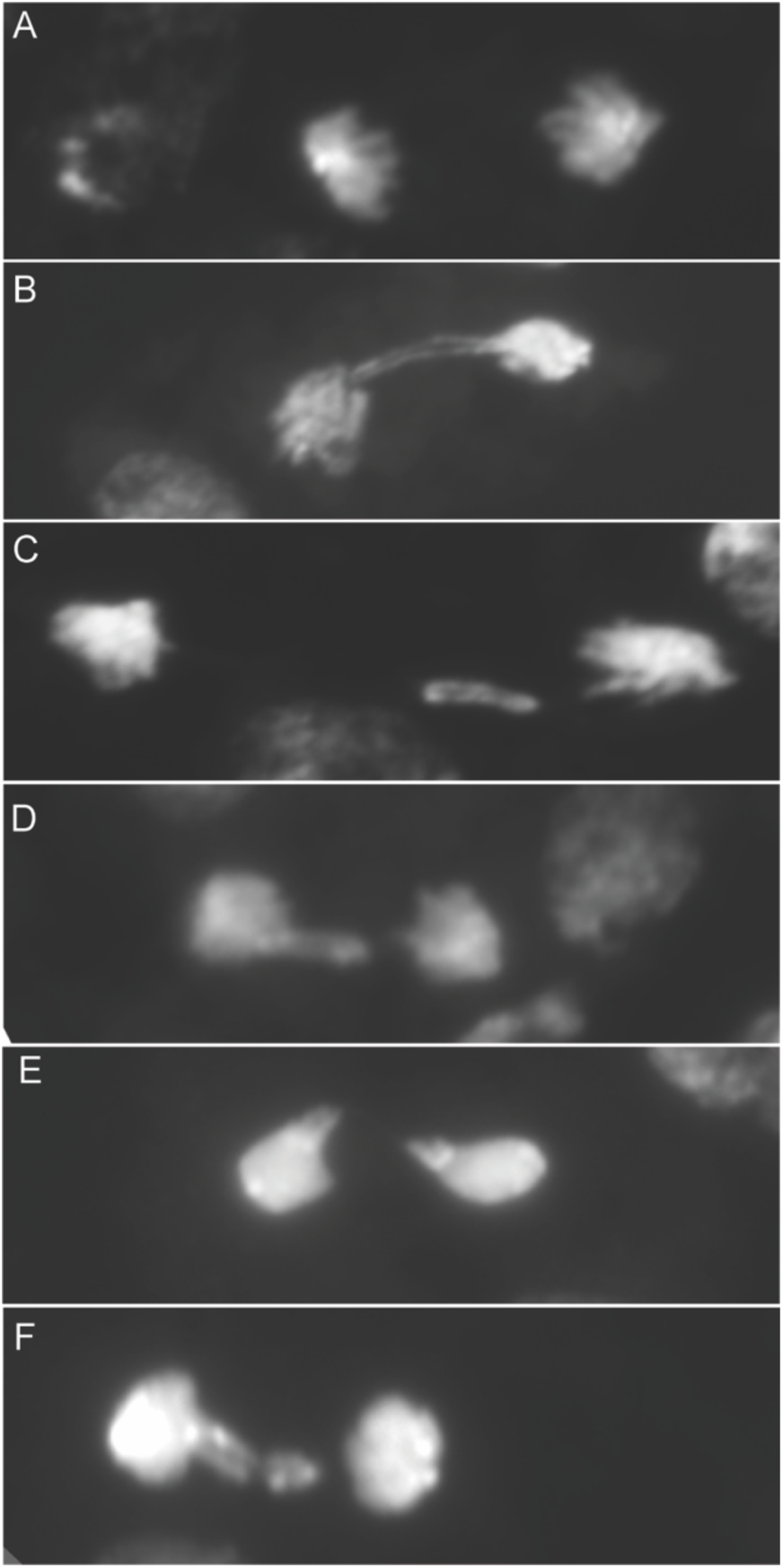
Ring dicentric bridges formed by heatshock-FLP in somatic neuroblasts (quantified in **Table 6**). A. a wildtype anaphase; B. a double dicentric bridge; C,D. lagging bridges; E,F. Broken double dicentric bridges

**Table 6:** Cytology of dicentric bridges in larval brains after FLP induction

| FLP | Ring | Bridges<br>(N) | Connected | Lagging* | Broken |
| --- | --- | --- | --- | --- | --- |
| 70FLP3F | *R(1;Y)6AX2* | 115 | 47% | 24.3% | 28.7% |
|  | *R(1;Y)11AX2* | 41 | 41.5% | 17% | 41.5% |
|  | *R(1)2* | 124 | 38.7% | 37.9% | 23.4% |
| 70FLP10 | *R(1;Y)6AX2* | 91 | 39.5 % | 25.3% | 35.2% |
|  | *R(1;Y)11AX2* | 134 | 51.5% | 16.4% | 32.1% |
|  | *R(1)2* | 141 | 48.2% | 30.5% | 21.3% |

\*Lagging describes bridges that have separated from one or both spindles

There is, however, another factor that might contribute to the differences in frequencies of linearized chromosomes in the germline and in the soma. It is notable that, in the somatic experiment, the frequency of linearized *R(1)2* chromosomes decreased at the 16-18 hr time point, while corresponding frequencies for *R(1;Y)6AX2* and *R(1;Y)11AX2* increased. Because *R(1)2* tends to break in euchromatin most often it may be that *R(1)2* is most likely to make linear products that are aneuploid, which are quickly eliminated. In contrast, most breakpoints of *R(1;Y)11AX2* were concentrated in a small region of heterochromatin, generating products would have no or minimal aneuploidy, perhaps allowing cells that receive them to survive longer. Although *R(1;Y)6AX2* and *R(1)2* both had frequent breaks in euchromatin, *R(1;Y)6AX2* exhibited more prominent hotspots in euchromatin than *R(1)2,* which might lead to more frequent coincidence of the two breaks and less frequent occurrence of aneuploid products, allowing such cells to survive longer than cells with broken *R(1)2* products. The depression in the appearance of broken *R(1)2* chromosomes in metaphases at 16-18 hrs after heat shock tends to support the view that many of the broken products are eliminated by a mechanism that more strongly affects *R(1)2*.

If *R(1)2* broken dicentrics are eliminated more frequently by cell death, then we might expect to see that directly reflected in an assay of cell death. Accordingly, we used Acridine Orange (AO) staining to examine cell death in wing discs from female larvae that carried one of the ring chromosomes and *70FLP10*. Larvae were heat shocked, and 24 hours later, wing discs were dissected from third instar larvae and stained with AO. There was a small amount of cell death in the absence of heat shock and there was a significant amount of cell death that occurred without FLP, simply as a result of heat shock. Cell death was greatly increased with heat shock induction of *70FLP* and consequent formation of dicentric chromosomes, but there was no significant difference in the occurrence of cell death between the three ring chromosomes (**Figure 10**). Although small differences in the frequencies of cell death are likely undetectable by this assay, these results suggest that a simple differential in the frequency of cell death is not sufficient to explain why linearized chromosomes are recovered at such different rates with the three tested rings.

**Figure 10:**
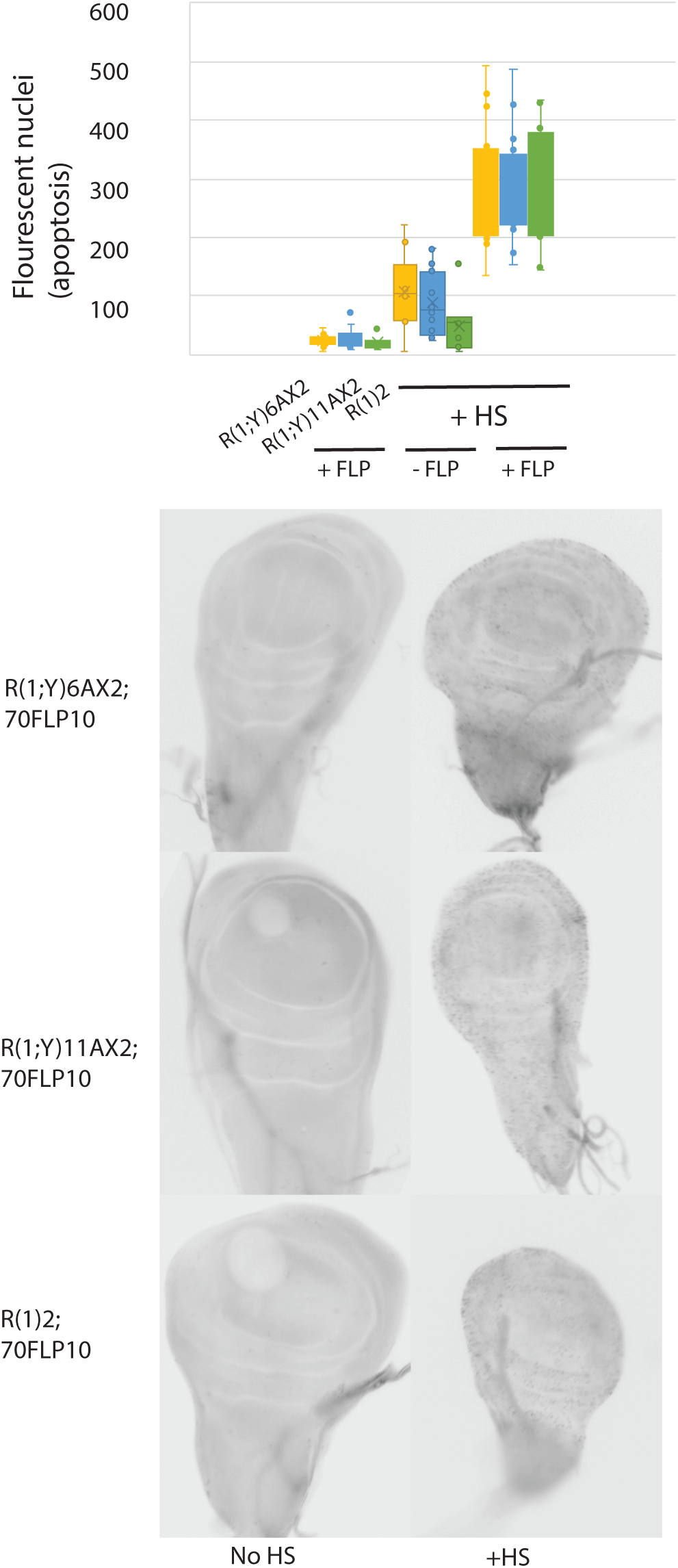
Detection of apoptosis in wing imaginal disks 24 hours after dicentric formation. Top: quantification of Acridine orange staining using ImageJ (Materials and Methods); Bottom: examples of the Acridine Orange staining viewed in wing imaginal disks

## Discussion

### What is a fragile site?

Fragile sites are, most simply, regions of chromosomes that are prone to breakage. In humans and other vertebrates the occurrence of fragile sites is linked with incomplete replication prior to mitosis. Under conditions that impair DNA polymerization, such as nucleotide starvation, chromosomal sites that replicate late appear to be especially susceptible to incomplete replication. These sites then appear as novel constrictions in metaphase chromosomes and may exhibit breakage, with spontaneous chromosome rearrangements often occurring at these sites (Arlt *et al*. 2006; Blumrich *et al*. 2011; Hussein *et al*. 2011; Fungtammasan *et al*. 2012). Underreplicated sites in *Drosophila* polytene chromosomes also exhibit fragility as they occasionally break during squashing (Bridges 1935; Lefevre 1976). The set of fragile sites known as Common Fragile Sites (CFS) are also well-conserved (Helmrich *et al*. 2006; Helmrich *et al*. 2007).

Our results show that *Drosophila* mitotic chromosomes have sites that are especially susceptible to breakage during dicentric chromosome segregation, meeting the most basic definition of fragile sites. However, unlike CFS in vertebrates, these sites are not conserved within *Drosophila melanogaster*: Chromosomes with different genealogical histories have different sets of dicentric breakage fragile sites. There is some coincidence with sites that are underreplicated in polytene chromosomes, but the correlation is incomplete, leading to the possibility that dicentric breakage hotspots represent a new type of fragile site. It is likely that other chromosomes in *Drosophila* possess similar breakage hotspots (Ahmad and Golic 1998). Maize and yeast chromosomes also exhibit hotspots of dicentric chromosome breakage (McClintock 1941; Pobiega and Marcand 2010; Song *et al*. 2013; Lopez *et al*. 2015), and possibly human chromosomes as well (Shimizu *et al*. 2005).

### What is responsible for breakage hotspots?

We previously considered the possibility that preferential sites of healing, rather than breakage, might account for the nonrandom distribution of ring opening sites. If chromosome breakage, by whatever means, occurs at random sites, but viable linear chromosomes are only recovered when breakpoints occur at preferred sites of healing, then the array of hotspots should not differ with different methods of breakage. As we reported previously, when ring chromosomes were opened by breakage with X-rays, the pattern of breakpoints was as we expected for random breaks — nearly all were in heterochromatin, presumably because breaks in heterochromatin need not be at the same site in each arm to generate a chromosome that is male-viable (Hill and Golic 2015).

To further test the underlying idea that X-rays induce random breaks, we examined existing data on *X* chromosome breakpoints of *X-*autosome translocations induced by ionizing radiation in *Drosophila melanogaster* (**Figure 11A**) (Bauer *et al*., 1938). We excluded division 20 from this analysis because those workers placed breakpoints in heterochromatin within division 20. The distribution of X-ray induced breakpoints was not significantly different than a Poisson distribution (assuming numbered divisions are equal in size; **Table 2** right, P = 0.7413). Furthermore, the distribution of breaks induced by γ-rays in *Drosophila heteroneura* also conforms to a Poisson distribution (Tonzetich *et al*. 1988). These findings strengthen the conclusion that the nonrandom distribution of breakpoints that we see following dicentric breakage is not a result of random breakage followed by preferential sites of healing. If it were, X-ray induced openings should show the same distribution, but they did not (Hill and Golic 2015).

**Figure 11:**
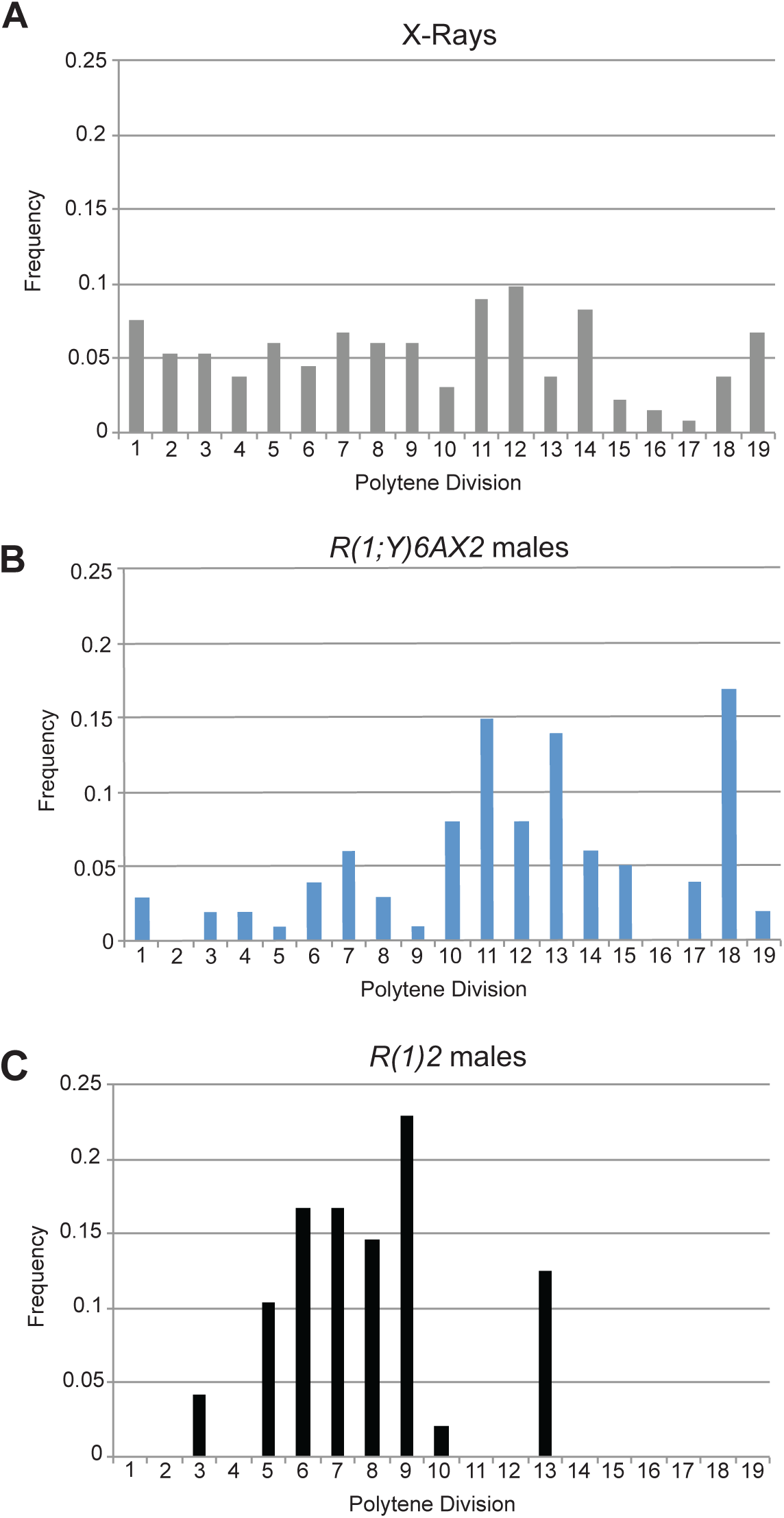
Distributions of euchromatic break locations and frequencies. A. X-ray induced rearrangements (from Bauer 1938); B. breakpoints of *R(1;Y)6AX2* openings recovered in males; C. breakpoints of *R(1)2* openings recovered in males. *R(1;Y)6AX2* and *R(1)2* tend to break in different halves of the euchromatin, to the right and left of division 10 respectively.

Another possible explanation for clustering might be that breaks occur randomly, but those occurring in areas of low gene density tend to be recovered because they are less likely to disrupt vital genes. Using coordinate data from CytoSearch, we found the overall gene density in *X* euchromatin is 115.7 genes/Mb (2661 genes 23MB). Our most extensive data set is for openings of *R(1;Y)*6AX2, and the gene density for breakage hotspots is 129.2 genes/Mb (divisions 11, 13, 18; 3.56 MB; 460 genes), and 121.2 genes/Mb (divisions 2, 5, 9, 16; 4.11 MB; 498 genes) for coldspots. If we assume that vital genes are distributed evenly throughout these regions (an assumption that cannot be currently validated) breakage hotspots cannot be explained by selection for breaks in regions of low gene density. This hypothesis also predicts that there should be similar hotspots for each of the three different rings and this is clearly not the case. Differential gene density does not account for breakage hotspots.

Another possible explanation for the hotspots is that cytokinesis is responsible for breakage. Perhaps the actin-myosin contractile ring provides the force necessary to break a stretched chromosome or facilitates the action of endonucleases that cut chromosome bridges (**Figure 12B**). If this were true, we expect that chromosome breakpoints would show a roughly Gaussian distribution around the midpoint of the bridges. In the case of *R(1;Y)6AX2,* if divisions 2-16 are considered by themselves, the breakpoint distribution somewhat approximates a normal distribution (P = 0.082).

**Figure 12:**
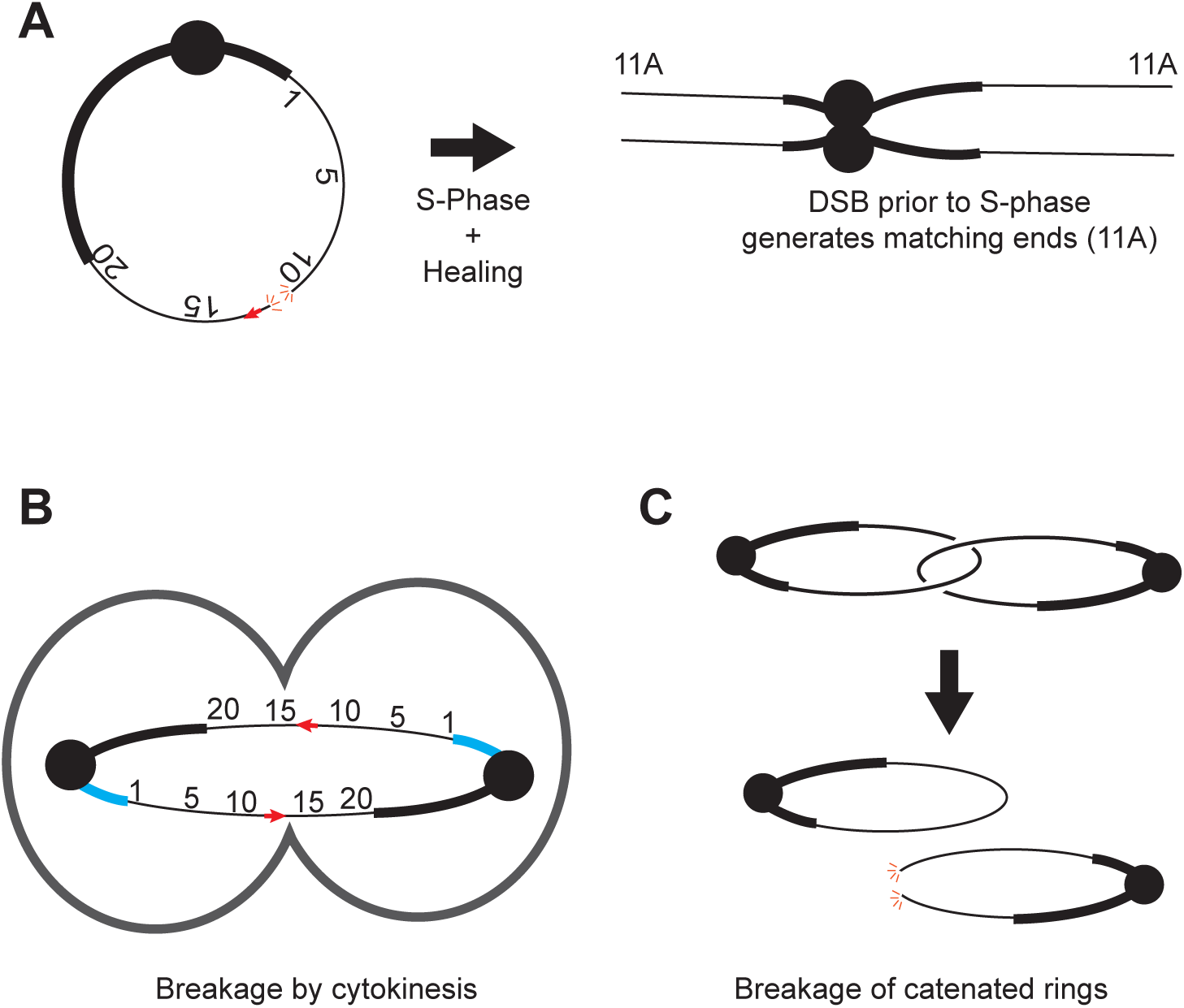
Potential explanations for breakage hotspots. A. Breakage and healing prior to replication would produce matching ends; B. Breakage by the cleavage furrow could generate breaks at approximately matching sites on the two bridges, the division plane is far from the small block of heterochromatin highlighted in blue; C. Catenated rings might be resolved by breakage of one ring, generating matched ends on the broken chromosome.

However, when the entire euchromatic portion of the *X* is considered, the breakpoints are not normally distributed (P < 0.005), no doubt due to the extreme hotspot in division 18. Additionally, the presence of hotspots in cytological divisions 11 and 13, but not 12, is difficult to explain under this hypothesis. Finally, some of the chromosomes with heterochromatin at each end cannot be readily explained by cytokinesis mediated cleavage. Especially in the case of breaks in the short heterochromatic arm, both breakpoints will be far from each other in the double dicentric bridges, and also far from the cleavage furrow.

A second prediction of the hypothesis that bridges are cleaved by cytokinesis is that the arms of linear derivatives should be of equal length. *R(1;Y)11AX2* derivatives provide an especially useful test of this prediction because more than half of the breaks were localized to a single region near the boundary between the heterochromatic long arm of the *Y* (*YL*) and X euchromatin. Accordingly, we measured both arms of six independent derivatives of *R(1;Y)11AX2* that terminated at this major hotspot. 44 metaphase images were examined and absolute arm lengths were determined. In 41/44 cases the predominantly euchromatic arm, consisting mostly of *X* euchromatin and a portion of *YS*, was longer than the predominantly heterochromatic arm, consisting mostly of *YL.* The ratio of arm lengths was 1.33 ± 0.08 (mean ± 2SE), arguing against the proposition that the bridges are cleaved by cytokinesis.

We also examined arm lengths for openings of *R(1;Y)6AX2* at its hotspots. For breaks in region 11 the ratio of the arm containing *YL* to the arm containing *YS* was 2.22 ± 0.25 (N=11); for region 13 the ratio was 1.29 ± 0.10 (N=16); for region 18 the ratio was 1.01 ± 0.08 (N=18). Only the hotspot in region 18 is compatible with a location in the center of the bridge. With *R(1)2* the location of the centromere is not apparent with DAPI staining so we did not determine arm length ratios for hotspots with this chromosome.

With ring chromosomes, there is also the possibility to form catenanes, either as the result of an even number of sister chromatid exchanges (Gatti *et al*. 1979), or by entanglement during replication, as happens with bacterial chromosomes. To successfully segregate these chromosomes, the interlock must be resolved. If topoisomerase fails to do the job, then breakage of one of the sister chromatids may occur instead (**Figure 12C**). This type of breakage cannot explain the large number of linearized chromosomes with duplications at their termini. Furthermore, such an event would most likely create only a single concentration of breakpoints directly opposite the centromere. Finally, if breakage of catenanes generated by replication were the source of linearized chromosomes then such chromosomes should be produced even without FLP-induced SCE. We tested this and found that linearized chromosomes were produced very infrequently without FLP (Table 1). This mechanism is insufficient to explain the breakage hotspots we detected.

### Why do fragile sites vary between chromosomes?

Although the euchromatin of the three rings is indistinguishable by polytene cytology the heterochromatin of each is morphologically unique, potentially accounting for some of the differences between these chromosomes. With *R(1;Y)11AX2,* breakage sites were located almost exclusively in heterochromatin — most near where the tip of *YL* is joined to *X* euchromatin. Owing to its method of construction *R(1;Y)11AX2* has *rDNA* repeats at this location (Golic and Golic 2011; confirmed by Ferree *et al*. 2014). The *rDNA* repeats are known to be unstable, and could be the basis for this particular hotspot (Lu *et al*. 2018; Ji *et al*. 2019). However, it is not clear why this *rDNA* cluster should be a greater breakage hotspot than the normal, and larger, *rDNA* locus embedded in *YS* on the same chromosome. It may be a position effect related to its unusual juxtaposition to euchromatin.

It is more difficult to devise a satisfactory explanation for the divergence of breakage hotspots within the euchromatin of *R(1;Y)6AX2* and *R(1)2*. One can imagine a number of reasons that a chromosome region might be prone to breakage under tension. In humans, CFS generally correspond to late-replicating regions and perhaps the dicentric breakage hotspots represent late-replicating, and occasionally unreplicated, regions. If a cell enters mitosis with unreplicated DNA the sister chromatids might still, with the aid of a variety of enzymes such as the MUS81-EME1 endonuclease (Naim *et al*. 2013; Ying *et al*. 2013; Calzetta *et al*. 2020), segregate normally and complete replication after mitosis. But, if this unreplicated DNA were found in a dicentric bridge it is easy to imagine that it would be weak point under tension. There is some overlap between the location of hotspots on each ring and sites of late/underreplication, but not all hotspots are explained by those studies (Zhimulev and Belyaeva 2003; Schwaiger *et al*. 2009; Belyaeva *et al*. 2012; Yarosh and Spradling 2014). Heterochromatin is known to be late-replicating and the *R(1;Y)11AX2* hotspots are all in heterochromatin. Polytene divisions 11 and 13 are known to be late-replicating in some cell types and are sites of frequent breakage in *R(1;Y)6AX2*. On the other hand, another late replicating region, 9A, is a breakage hotspot in *R(1)2* but a coldspot in *R(1;Y)6AX2*. Since the basis of replication origin choice is not well understood, and replication timing can differ between cell types and cell lines (Schwaiger *et al*. 2009; Hua and Orr-Weaver 2017) it is quite conceivable that preferred origins, or regions devoid of origins, might vary between strains. However, if dicentric breakage hotspots also correspond to late-replicating regions it is surprising that we saw no effect of aphidicolin on breakage frequency or hotspot distribution.

In human cells CFS also tend to be associated with large transcribed genes (Smith *et al*. 2006; Bosco *et al*. 2010). Dicentric breakage hotspots might arise because transcription has occurred, or is still occurring, as cells enter the mitotic phase of the cell cycle and the presence of an open helix might generate a site susceptible to breakage. Conflicts between transcription and replication have also been linked to CFS in humans (Tuduri *et al*. 2009; Helmrich et al. 2011). Transcriptional activity might also lead to a regional difference in chromatin packaging or chromosome condensation in mitosis, resulting in localized decondensation as tension is applied during anaphase. Such a region might be more prone to breakage by tension, or might instigate cleavage by specialized enzymes, *e.g.* the TREX1 endonuclease (Maciejowski *et al*. 2015). This explanation would also require that different ring chromosomes differ in their pattern of transcription in male germline stem cells.

Another possibility is that breakage might be connected to sites of transposon insertion in these chromosomes. This idea is appealing because transposons are numerous, are distributed nonrandomly throughout the genome (Kaminker *et al*. 2002), and their mobility provides an explanation for differing distributions between strains that have been separated for many generations. There are several ways in which transposons might provide the basis for breakage hotspots. First, as discussed above, breakage hotspots might be transcription-based. Transposons might be transcribed late in the cell cycle, perhaps as a strategy to evade piRNA-mediated silencing (Czech *et al*. 2018), or even during mitosis as a consequence of chromatin stretching. Alternatively, if a mechanism to transcriptionally silence transposons via the piRNA pathway exists in the male germline, as it does in the female germline (Le Thomas *et al*. 2013), the altered chromatin configuration associated with silencing might generate breakage hotspots. Breakage hotspots could also result from occasional transposon mobilization. A double strand break generated by mobilization of a DNA transposon during mitosis would cleave a dicentric bridge, allowing completion of cell division. Alternatively, attempted insertion of a transposon, perhaps cued by stretched chromatin during anaphase, could also produce a break. There is at least one example of a preference for viral integration at CFS in humans (Popescu et al. 1990; Wilke et al. 1996; Thorland et al. 2000; Gao et al. 2017).

One final possibility that should be considered is that there are multiple causes of breakage hotspots and no single explanation will suffice to explain all. For instance, *R(1;Y)6AX2* has two hotspots in late replicating regions (#11 & #13) and the third is in the middle of the dicentric bridge (#18). It might be possible to adduce evidence for or against these various hypotheses by extensive sequencing of the parent ring chromosome and their linear derivatives.

### What causes breakage of a dicentric bridge?

Barbara McClintock was the first to show, in maize, that dicentric bridges can break. The ensuing consequences include the well described Breakage-Fusion-Bridge cycle and aneuploidy (McClintock 1941). On the other hand, Sturtevant and Beadle (1936) deduced that meiotic dicentric bridges did not break in *Drosophila melanogaster*. In comparing the divergent fates of bridges in plants and animals it was suspected that dicentric bridges might require cleavage by the plant cell wall during cytokinesis (McClintock 1938), accounting for a failure of breakage in animal cells. Although dicentric bridges may not always break in animal cells (Pampalona *et al*. 2016), dicentric breakage has been observed in many organisms besides plants, including *Saccharomyces cerevisiae*, *Drosophila melanogaster*, and vertebrates. (Ahmad and Golic 1999; Zhou and Elledge 2000; Thrower and Bloom 2001; Lo *et al*. 2002; Sancar *et al*. 2004; Titen and Golic 2008; Maciejowski *et al*. 2015). Since dicentric chromosomes can break in organisms without cell walls, the inevitable question is, what force is responsible for breakage? (See Guérin and Marcand 2022 for a recent review of this topic.)

An obvious possibility is that the tension applied to a dicentric chromatin bridge during anaphase is the cause of breakage. This raises the question of whether a mitotic spindle is capable of exerting sufficient force to break a dsDNA molecule. In *S. cerevisiae*, where dicentric chromosomes have been produced by chromosomal integration of the 2µ plasmid, by activation of a conditional centromere, or by fusion of dysfunctional telomeres (Falco *et al*. 1982; Hill and Bloom 1989; Brock and Bloom 1994; Ferreira and Cooper 2001; Pardo and Marcand 2005) it is unlikely that the spindle can break a chromosome. Each kinetochore binds one microtubule which has an estimated maximum pulling force of 30-65 pN (Winey *et al*. 1995; Grishchuk *et al*. 2005), though the actual tension on a yeast chromosome during normal segregation is about an order of magnitude less (Chacón *et al*. 2014). While this is enough force to physically segregate sister chromatids in the viscous cytoplasm, it is likely not enough to break a dsDNA molecule: the force required to break DNA has been measured at between ∼270-470 pN (Harrington and Zimm 1965; Bensimon *et al*. 1995). In *S. cerevisiae,* cytokinesis is generally required to break dicentric chromosomes (Lopez *et al*. 2015). It is not known whether the bridge is cut by physical forces at the cleavage furrow or whether nucleases assist in the process. In human cells the TREX1 endonuclease contributes to dicentric bridge breakage during cytokinesis (Maciejowski *et al*. 2015), though cytokinesis may not always be required for bridges to break (Shimizu *et al*. 2005).

Higher eukaryotes have larger kinetochores that attach multiple microtubules and it is possible that spindle forces are capable of breaking a dicentric chromosome. Calibrated microneedles were used to determine the force produced on a chromosome by the spindle in grasshopper spermatocytes: approximately 700 pN was required to stall poleward movement of a chromosome during division (Nicklas 1983). In *Drosophila* S2 cells spindle force was measured using a FRET tension sensor placed within the Cenp-C protein, leading to the conclusion that the 11 MTs that typically attach to each kinetochore generate 135-680 pN of pulling force on the kinetochore (Ye *et al*. 2016).

In order for spindle tension to cause a chromatin bridge to break, the tension must be applied directly to the DNA. Because anaphase chromosomes are highly folded structures, one might imagine that considerable stretching may occur before the dsDNA molecule experiences sufficient force to cause breakage. However, if a stretched chromosome first unraveled at one or a few locations, it could generate a region where breakage might ultimately occur. CFS in human cells are known to experience abnormal condensation during mitosis (Boteva *et al*. 2020), which, in a dicentric chromosome, might make those sites especially susceptible to breakage. Unravelling of a chromatin fiber can happen with only 5-6 pN of force, but becomes irreversible after ∼20pN, presumed to reflect the loss of nucleosomes (Cui and Bustamante 2000). As tension continues to rise, dsDNA denatures to form single strands of DNA. Strand separation can occur *in vitro* with 35-300 pN force (depending on A/T content) and ∼2x stretching distance (Noy *et al*. 1997; Clausen-Schaumann *et al*. 2007). Finally, the tension reaches a threshold and covalent bond breakage must occur somewhere within the DNA backbone of both ssDNA strands, requiring <500 pN. Together, these results suggest that, in higher eukaryotes, the spindle may be capable of producing sufficient force to break DNA.

### Centromere size, strength and chromosome breakage

The question of whether dicentric bridges break under tension was addressed by Sturtevant and Beadle nearly a century ago (1936; recently reviewed by Hawley and Ganetzky 2016). By examining outcomes of dicentric bridges formed in *Drosophila* female meiosis, where cytokinesis does not occur, they concluded that dicentric bridges with two normal *X* centromeres do not break. Subsequently, Novitski examined the fate of dicentric chromosomes with *Y* chromosome centromeres and concluded that those dicentric bridges did break. Novitski proposed that the *X* chromosome centromere was weak and the *Y* centromere was strong, accounting for bridges remaining intact in the former case and breaking in the latter. Associated heterochromatin might also contribute to these differences in strength (Lindsley and Novitski 1958).

In the work reported here, a correlation emerged between the origin of the centromere plus its surrounding heterochromatin and the frequency of linear chromosome recovery. In the germline, linear forms of *R(1;Y)11AX2*, with a *Y* centromere and substantial surrounding heterochromatin, were recovered about twice as frequently as *R(1;Y)6AX2,* which also carried a *Y* centromere but with less heterochromatin, and about 10 times more frequently than *R(1)2,* with its *X* centromere (**Table 1**). In the soma, linearized *R(1;Y)11AX2* were also seen more frequently than the other rings, and at the later time point linearized *R(1;Y)6AX2* chromosomes were seen more frequently than linearized *R(1)2* (**Figure 8**). When we examined the production of anaphase bridges, another relationship emerged: *R(1)2* bridges lagged during division more often than bridges produced with the other rings, suggesting that one or both kinetochores had become detached from the spindle (**Table 6**).

Our observations support the principle of the strong/weak centromere hypothesis put forth by Novitski: that the spindle pulls the *Y* centromere more forcefully than the *X* centromere (Novitski 1952; Novitski 1955). In further support of this hypothesis, the *Drosophila Y* chromosome kinetochore is approximately twice as large as the *X* kinetochore (Lin *et al*. 1981; Raychaudhuri *et al*. 2012). We also note that S2 cells, where Ye *et al*. measured force on the kinetochore (2016), do not carry a *Y* chromosome (Zhang *et al*. 2010) and their measurements would not account for the larger *Y* centromere. Considering the large variation in measurements of forces required to break DNA and forces exerted by the spindle on segregating chromosomes, it is easy to imagine that the *X* and *Y* centromeres might differ substantially in their attachment to spindle microtubules, and in such a way that the *X* centromere does not typically generate the force needed to break a dicentric bridge, but the *Y* centromere does.

The question of how dicentric chromosomes break, and where they break, has broad human health implications and evolutionary significance. Dicentric chromosomes are a frequent occurrence in cancer cells (Cleal and Baird 2020; Dewhurst 2020). Breakage hotspots will influence whether cells experience altered dosage of genes that regulate the growth of cancer cells (LeTallec *et al*. 2013; Glover *et al*. 2017; Simpson *et al*. 2021). Chromosome breakage can also lead to rearrangements that alter expression of genes that promote or impede carcinogenesis (Zheng 2013). Chromosome rearrangements, such as chromosome fusion and fission events also accompany evolutionary divergence of species (White 1973; Mayrose and Lysak 2021). Understanding the mechanisms that produce dicentric breakage hotspots, and the degree to which that mechanism converges or diverges from mechanisms that produce CFS will be important to fully understand how genome and karyotype evolution occurs, both over the time scale of tumor growth and the time scale of evolution.

## Funding

This work was supported by grant R35 GM136389 from the National Institutes of Health.

## Notes

### Competing Interest Statement

The authors have declared no competing interest.

