## Supplemental Figure S5 for "Dicentric chromosome breakage in *Drosophila melanogaster* is influenced by pericentric heterochromatin and reveals a novel class of fragile site"

Worksheet 1: 6AX2 - 19 bins

Chi-Square Goodness-of-Fit Test for Observed Counts in Variable: C2

**Observed and Expected Counts**

| **Category** | **Observed** | **Test Proportion** | **Expected** | **Contribution to Chi-Square** |
| --- | --- | --- | --- | --- |
| 1 | 9 | 0.22346 | 4.24574 | 5.32369 |
| 2 | 2 | 0.16347 | 3.10593 | 0.39379 |
| 3 | 1 | 0.17380 | 3.30220 | 1.60503 |
| 4 | 2 | 0.15398 | 2.92562 | 0.29285 |
| 5 | 0 | 0.11693 | 2.22167 | 2.22167 |
| 6 | 5 | 0.16836 | 3.19884 | 1.01417 |

6 (100.00%) of the expected counts are less than 5.

**Chi-Square Test**

| **N** | **DF** | **Chi-Sq** | **P-Value** |
| --- | --- | --- | --- |
| 19 | 5 | 10.8512 | 0.054 |

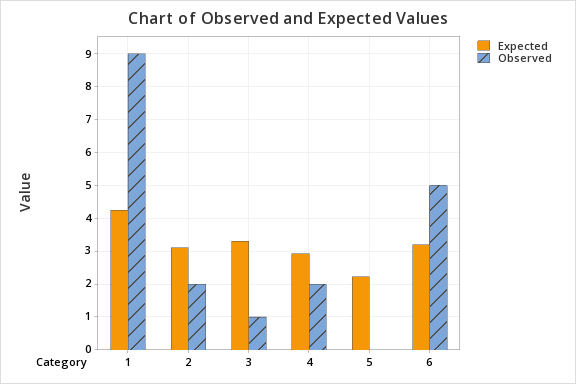

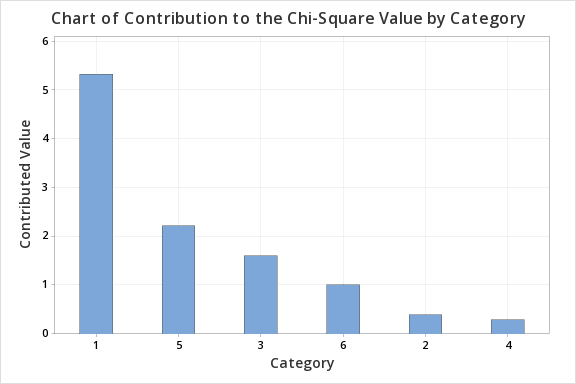

Worksheet 2 : 6AX2 - 22 bins

Chi-Square Goodness-of-Fit Test for Observed Counts in Variable: C2

**Observed and Expected Counts**

| **Category** | **Observed** | **Test Proportion** | **Expected** | **Contribution to Chi-Square** |
| --- | --- | --- | --- | --- |
| 1 | 9 | 0.15254 | 3.35588 | 9.49262 |
| 2 | 3 | 0.15752 | 3.46544 | 0.06251 |
| 3 | 3 | 0.18499 | 4.06978 | 0.28120 |
| 4 | 0 | 0.17380 | 3.82360 | 3.82360 |
| 5 | 2 | 0.13608 | 2.99376 | 0.32987 |
| 6 | 1 | 0.09132 | 2.00904 | 0.50679 |
| 7 | 4 | 0.10375 | 2.28250 | 1.29236 |

7 (100.00%) of the expected counts are less than 5.

**Chi-Square Test**

| **N** | **DF** | **Chi-Sq** | **P-Value** |
| --- | --- | --- | --- |
| 22 | 6 | 15.7890 | 0.015 |

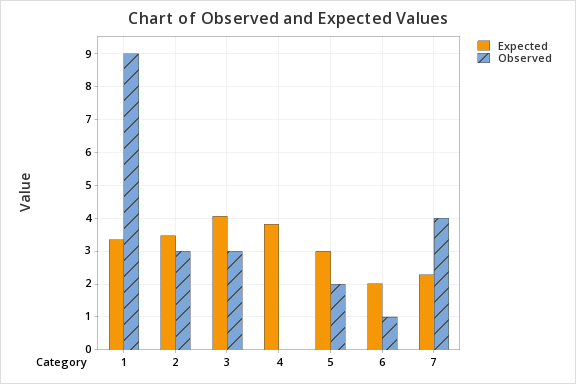

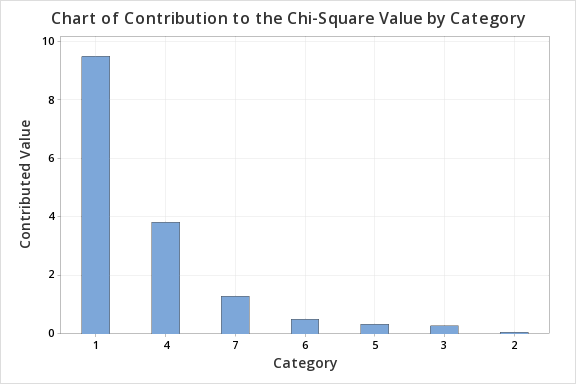

Worksheet 3 : X-Ray breakage 19 bins

#### Chi-Square Goodness-of-Fit Test for Observed Counts in Variable: C3

### Observed and Expected Counts

| **Category** | **Observed** | **Test Proportion** | **Expected** | **Contribution to Chi-Square** |
| --- | --- | --- | --- | --- |
| 1 | 4 | 0.17297 | 3.28643 | 0.15493 |
| 2 | 3 | 0.12771 | 2.42649 | 0.13555 |
| 3 | 1 | 0.14900 | 2.83100 | 1.18423 |
| 4 | 2 | 0.14900 | 2.83100 | 0.24393 |
| 5 | 3 | 0.13037 | 2.47703 | 0.11041 |
| 6 | 6 | 0.27095 | 5.14805 | 0.14099 |

5 (83.33%) of the expected counts are less than 5.

### Chi-Square Test

| **N** | **DF** | **Chi-Sq** | **P-Value** |
| --- | --- | --- | --- |
| 19 | 5 | 1.97005 | 0.853 |

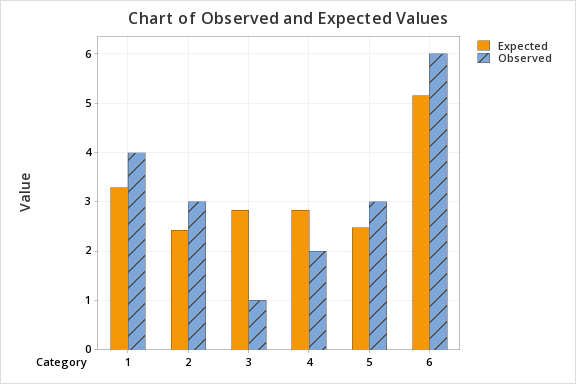

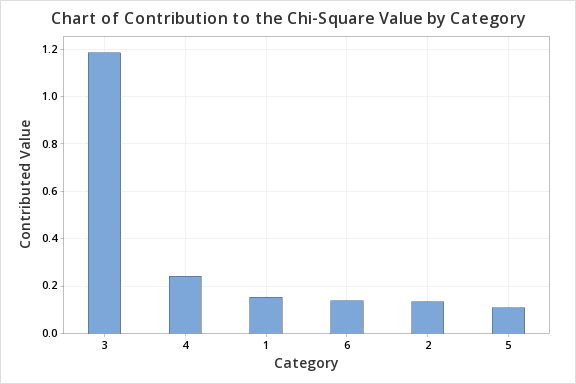
