## Supplementary figures and images for "Dicentric chromosome breakage in *Drosophila melanogaster* is influenced by pericentric heterochromatin and reveals a novel class of fragile site"

### Supplemental figure S4

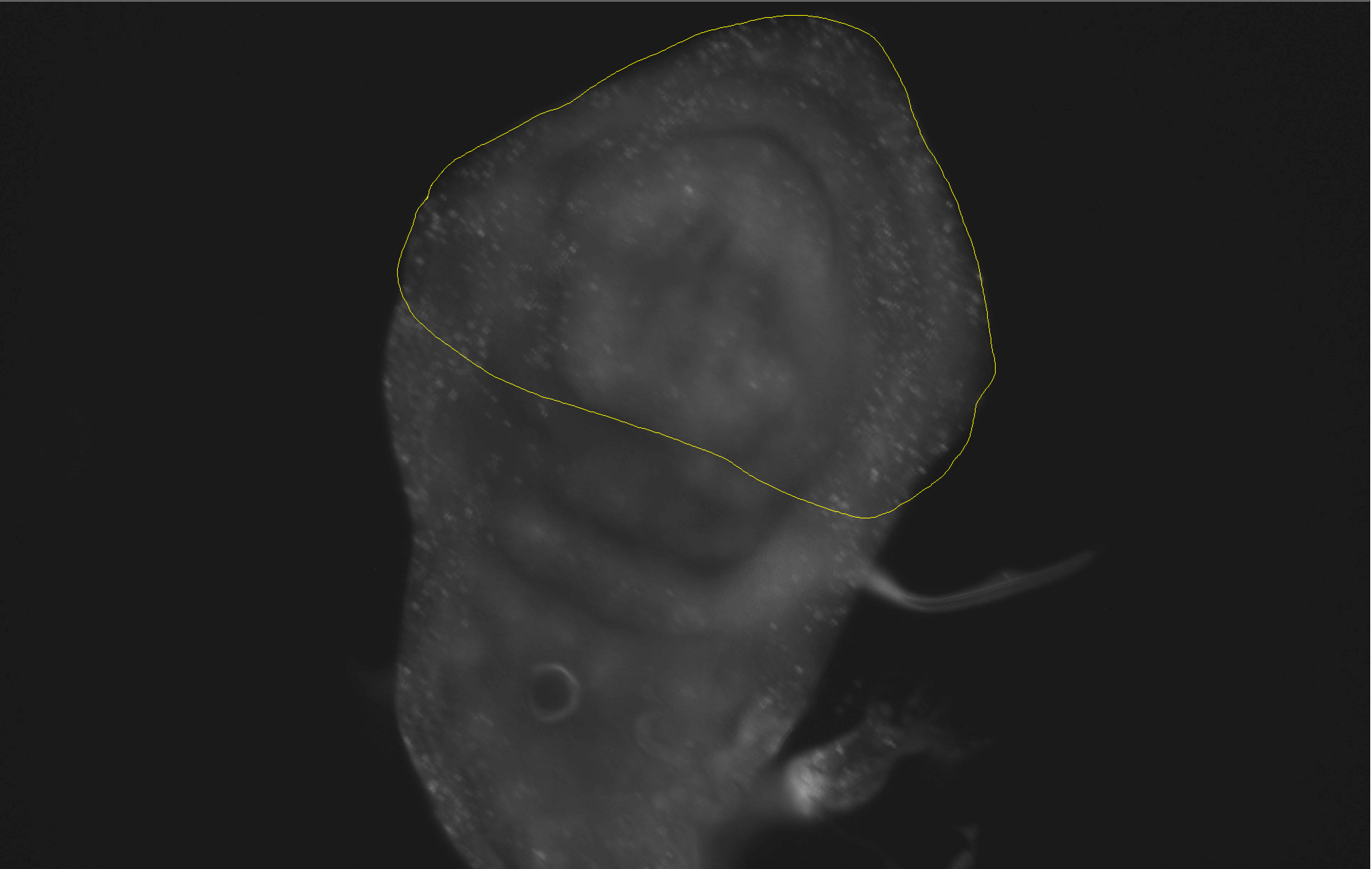
